# Transgenerational fitness effects of lifespan extension by dietary restriction in *Caenorhabditis elegans*

**DOI:** 10.1101/2020.06.24.168922

**Authors:** Edward R. Ivimey-Cook, Kris Sales, Hanne Carlsson, Simone Immler, Tracey Chapman, Alexei A. Maklakov

## Abstract

Dietary restriction increases lifespan in a broad variety of organisms and improves health in humans. However, long-term transgenerational consequences of dietary interventions are poorly understood. Here we investigated the effect of dietary restriction by temporary fasting (TF) on mortality risk, age-specific reproduction and fitness across three generations of descendants in *C. elegans*. We show that while TF robustly reduces mortality risk and improves late-life reproduction in the parental generation (P_0_), it has a wide range of both positive and deleterious effects on future generations (F_1_-F_3_). Remarkably, great-grandparental exposure to TF in early-life reduces fitness and increases mortality risk of F_3_ descendants to such an extent that TF no longer promotes a lifespan extension. These findings reveal that transgenerational trade-offs accompany the instant benefits of dietary restriction underscoring the need to consider fitness of future generations in pursuit of healthy ageing.

## Introduction

Dietary restriction (DR), a reduction in nutrient intake without malnutrition, is an environmental intervention that robustly extends lifespan and/or improves health across a broad cross-taxonomic variety of organisms from yeast to mice to primates (Weindruch *et al.*, 1989; Mair & Dillin, 2008; Nakagawa *et al.*, 2012; Fontana & Partridge, 2015; Mattison *et al.*, 2017). However, DR commonly reduces reproduction and long-term DR can be difficult to sustain in humans (Partridge *et al.*, 2005; Pifferi *et al.*, 2019). These considerations led to the development of alternative approaches such as DR mimetics and less demanding DR regimes such as different forms of temporary fasting (TF) (Lee & Longo, 2011; Selman, 2014). Temporary fasting, including intermittent fasting, has been shown to increase lifespan in model organisms (Mattson & Wan, 2005; Kaeberlein *et al.*, 2006; Lee *et al.*, 2006; Honjoh *et al.*, 2009; Uno *et al.*, 2013) and improve health in humans (Brandhorst et al. 2015), and some forms of TF are currently being investigated for their potential to speed up patient recovery after surgery and chemotherapy (Di Biase et al. 2016). Nevertheless, we know very little about the potential effects of DR, in whichever form it is implemented, on the fitness of offspring and even less so about the transgenerational effects of DR on the fitness of more distant descendants.

Despite the positive effects of DR, several studies show that it can also be costly. For example, a recent study in *Drosophila melanogaster* fruit flies reported reduced survival and fecundity in individuals returned to a standard diet after a period of fasting, suggesting a hidden cost of improved survival under DR (McCracken *et al.*, 2020). However the data from model organisms is currently inconclusive, because other studies in *D. melanogaster* did not report such effects (Mair *et al.*, 2003; Fricke *et al.*, 2008), while a study in *C. remanei* nematodes suggests that DR implemented through TF improves fitness upon return to normal feeding conditions (Mautz *et al.*, 2020). Nevertheless, there are good reasons to believe that DR occurring in the parents may affect offspring health and lifespan and that these effects will depend on the environmental conditions encountered by the offspring. Anticipatory parental effects may improve offspring performance when offspring themselves are raised under DR (Hibshman *et al.*, 2016) but may result in reduced fitness when offspring are raised in a standard environment (Harvey & Orbidans, 2011; Rando & Simmons, 2015). The detrimental effects of such environmental mismatches between parents and their offspring have been shown previously (Burgess & Marshall, 2014) but are rarely investigated as a potential fitness cost of DR-mediated lifespan extension. Nevertheless, while DR by TF increases fitness of ageing parents in *C. remanei* nematodes, it reduces fitness of their offspring in *ad libitum* food conditions (Mautz *et al.*, 2020).

Recently, there has been a surge of interest in transgenerational effects where parental condition affects the health and fitness of not only offspring and grand-offspring but also of more distant descendants. Such effects, if common, may have a profound influence on major evolutionary processes (Bonduriansky & Day, 2019) and will likely have important implications for translational research (Anway *et al.*, 2005). Specifically, transgenerational trade-offs between parental fitness and fitness of distant descendants may constitute an obstacle for research programmes aimed at harnessing the power of DR for life- and health-span extension (Maklakov & Immler, 2016). Alternatively, transgenerational transfer of the desired phenotype may be seen as an additional benefit. Some of the most spectacular examples of transgenerational effects come from recent work on *C. elegans* nematodes (Greer *et al.*, 2011; Rechavi, 2014; Rechavi *et al.*, 2014; Rechavi & Lev, 2017; Moore *et al.*, 2019; Burton *et al.*, 2020). Moreover, research suggests that larval starvation in ancestors results in transgenerational inheritance leading to increased lifespan of F_3_ offspring (Rechavi *et al.*, 2014). Thus, the combination of novel theoretical considerations and recent empirical discoveries calls for the investigation of transgenerational effects of ancestral DR on lifespan and fitness of descendants.

There are two main evolutionary models that explain the life-extending effect of DR despite the overall reduction in resources. The resource allocation model suggests that animals experiencing food shortage will temporarily switch their metabolism from reproduction to somatic maintenance in order to increase their chances of survival until resources will become plentiful again (Shanley & Kirkwood, 2000). An important extension of this model is the consideration of direct negative effects of reproduction on survival. Thus, by reducing reproduction in favour of survival, the animals both have relatively more resources for somatic maintenance and repair and suffer less damage from reproduction (Shanley & Kirkwood, 2000). However, a recent model suggested that DR-mediated lifespan extension may be an unselected by-product of increased autophagy with the goal of maximising reproduction using internal resources under the conditions when external resources are limited (Adler & Bonduriansky, 2014). Interestingly, recent studies in fruit flies and nematodes suggest that DR response is under neuronal control and can be manipulated by providing animals with food odour alone (Apfeld & Kenyon, 1999; Alcedo & Kenyon, 2004; Libert *et al.*, 2007; Smith *et al.*, 2008). The smell of food is sufficient to reduce the benefits of DR suggesting that animal physiology is switched from a self-preservation mode to reproduction mode. This technique provides us with an elegant research tool to test whether animals supress or maximise their reproduction under DR but so far there have been no studies of the transgenerational fitness consequences of such treatments.

Here we focused on addressing the following unresolved questions: 1) How does TF affect mortality risk and reproductive ageing once the animals return to their standard food regime?; 2) How do offspring of fasting parents perform in matching and mis-matching environments?; 3) Do transgenerational effects of ancestral fasting shape mortality risk and reproductive ageing of distant descendants?; and 4) Does reduced reproduction under DR represent a decision-making strategy? We use *C. elegans* nematode worms, which are an established model for the study of both DR and transgenerational effects, to investigate how two-day bacterial deprivation in early adulthood affects mortality risk and age-specific reproduction in ancestors and their descendants over a total of four generations. Our results reveal strong and previously unknown transgenerational costs of DR response and demonstrate that organisms supress their own reproduction under DR. We use the results of this study to make major inferences about the adaptive nature of DR response and suggest that transgenerational costs of DR can be sufficiently severe to be considered in any applied programme aimed at lifespan extension via reduced nutrient intake.

## Materials and Methods

### Strains

*Caenorhabditis elegans* nematodes of the Bristol N2 wild-type strain from the *Caenorhabditis* Genetics Center were used in all assays. Populations were thawed from −80°C and underwent bleaching and egg-lays prior to the start of the experiment in order to synchronise the developmental age of the P_0_ individuals. Populations were then kept in climate chambers set to 20°C, 60%RH and continual darkness. Worms were maintained on standard Nematode Growth Medium (NGM) agar in Petri dishes. These were 90mm wide for population maintenance and 35mm for individual culture. The NGM agar contained a fungicide (100 μg/ml Nystatin) and antibiotics (100 μg/ml Ampicillin and 100 μg/ml Streptomycin) to prevent infection. In all cases, nematodes were fed using antibiotic resistant *Escherichia coli* OP50-1 (pUC4K), from J. Ewbank at the Centre d’Immunologie de Marseille-Luminy, France. The OP50-1 (pUC4K) *E. coli* were thawed from −80°C, streaked on LB plates containing Ampicillin (100 μg/ml) and Streptomycin (100 μg/ml), incubated at 38°C for 16 ± 1 hours and kept at 6°C. A single bacterial colony was inoculated per 40 ml of LB broth containing the same antibiotic concentrations then incubated at 37°C for 16 ± 1 hours. The *E. coli* solution was then pipetted onto NGM agar Petri dishes and incubated at 20°C for 18 ± 6 hours to grow *ad libitum* bacterial lawns prior to use. The time from bacterial thawing to use was within a month. NGM agar plates were used within a few weeks (max = 4 weeks) of preparation.

### Experimental set-up

The transgenerational assay required the use of four different small plates (35mm), one for each of the 4 P0 treatments (Fig 1). (i) *Ad libitum* (AL) control plates consisted of standard NGM agar plates topped with an *E. coli* lawn. (ii) TF plates contained no peptone within the NGM agar and were not seeded with *E. coli*, which restricted the potential growth of a substitute food source. (iii) Food odour (FO) plates were designed so worms could detect the presence of an *E. coli* food source but were unable to access it. They consisted of a base NGM agar layer seeded with *E. coli* then incubated overnight. A second NGM agar layer, without peptone and without bacterial seeding, was then added. (iv) The fourth treatment (FO+AL) was a positive control plate, combining the food odour treatment with *ad libitum*, by layering a FO plate with a top agar layer containing peptone and bacterial seeding. The time allowed for bacterial growth between treatments was standardised. Two days prior to use, the bottom layers for the FO and FO+AL positive control treatments were incubated. All plates were incubated for one day in order to grow the upper seeding and to standardise incubation time across all four treatments.

**Fig 1.**
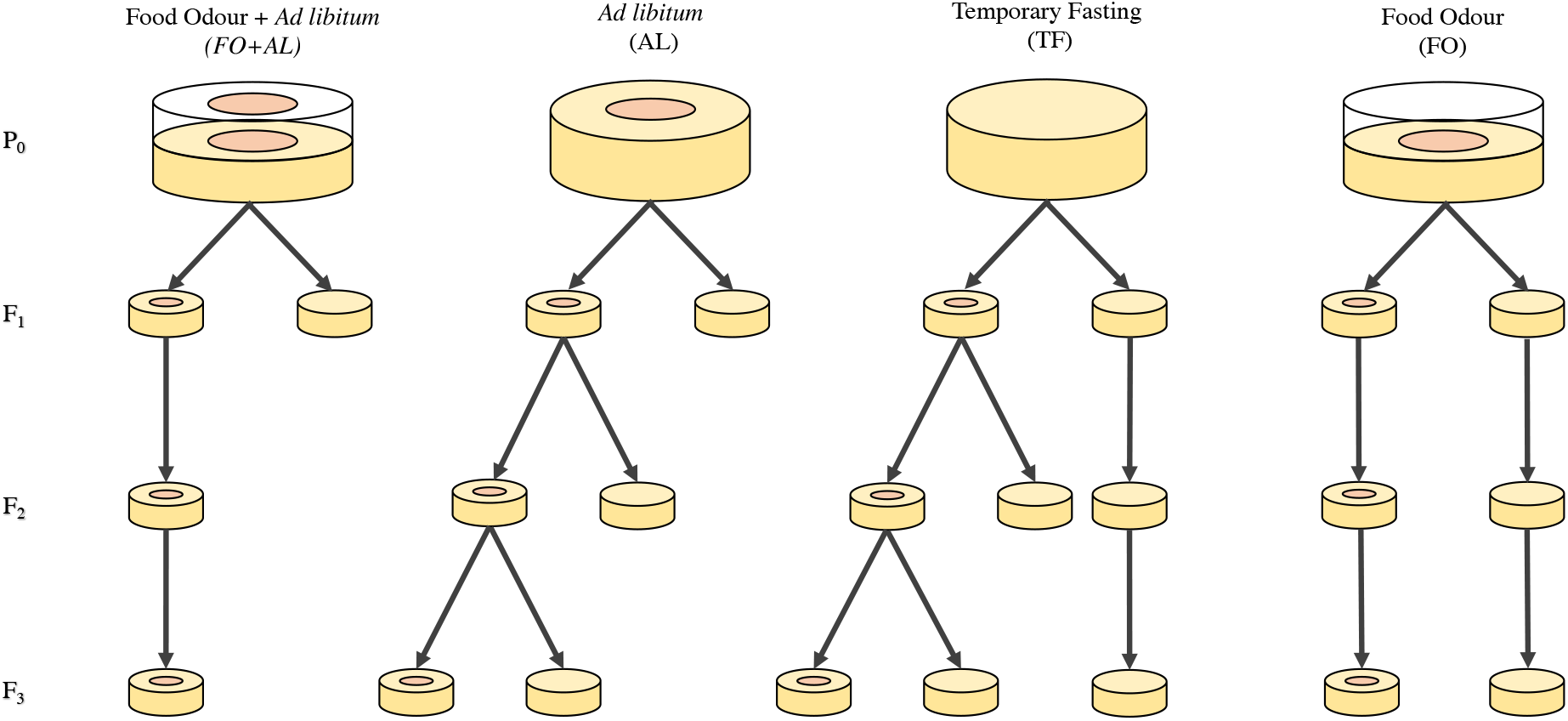
A visual representation of the dietary lineages studied (from P_0_ – F_3_). Not all dietary lineages were continued until the F_3_ generation (discussed below).

### Transgenerational lifespan and fitness assays

In order to study the transgenerational effects of P0 diet treatments, individual late L4 stage nematodes were randomly exposed to one of four dietary treatments (see above; sample sizes varied for each treatment owing to disproportionate day one and two mortality of TF individuals (*n* = 110 for AL/FO+AL, *n* = 191 for FO, *n* = 259 for TF). They remained exposed to these dietary conditions for two days prior to transfer onto standard *ad libitum* control plates seeded with *E. coli.* The presence or absence of food led to corresponding shifts in reproductive schedule (See Results). Thus, in order to generate sufficient progeny for future generations (F_1_-F_3_), we took offspring from different days for each of the four dietary treatments. For those that had access to food, the AL and FO+AL treatments, subsequent generations were generated from larvae taken from two-day old parents. For those that had restricted access to food, the TF and FO treatments, subsequent generations were generated from larvae taken from four-day old parents. The different choice of days allowed us to obtain sufficient offspring from all treatments and ensure that larvae were raised in fully fed environments across all treatments. To generate successive generations, eggs were allowed to hatch and develop for two days upon which two surviving larvae from each parent were randomly allocated into either AL or TF conditions (in some cases, this number was increased to compensate for higher than predicted loss of individuals; Fig. 3A-D and Fig. 5A-D). This continued for two further generations (until F_3_, see Fig. 1). In order to ensure a sufficiently large sample size across treatments and also due to lack of *a priori* expectations and the need to focus our tests around the questions above, certain lineages were only continued until F_1_ (see Fig. 1).

**Fig. 2.**
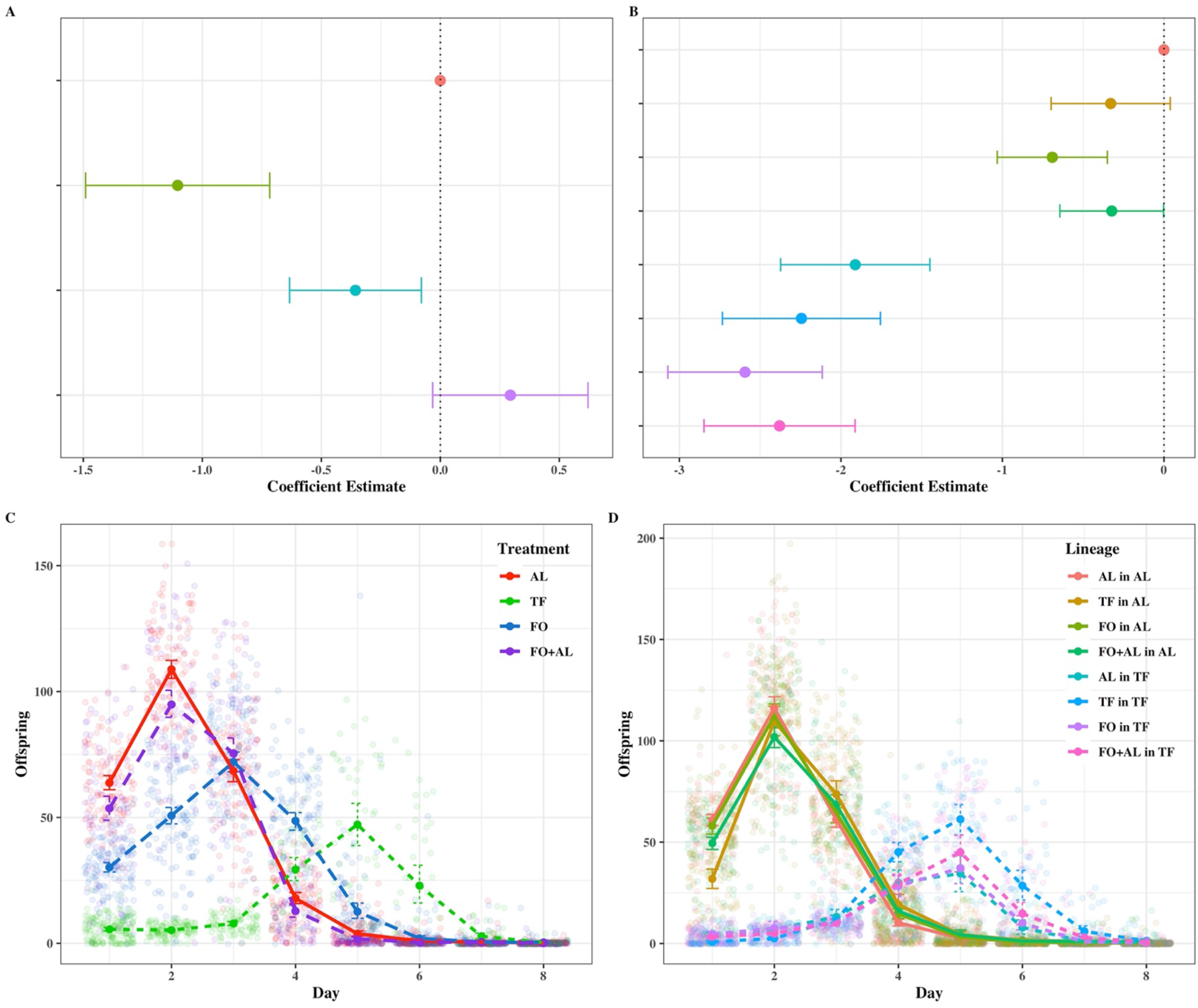
**A-D** Plots of survival (A-B) and reproduction (C-D) for P_0_ (A, C) and F_1_ (B, D). Colours represent differing dietary treatments for each generation. For Graph D the lineages are given as P_0_ in F_1_. A-B) Points represent coefficients from a mixed effects cox model with 95% confidence intervals; C-D) Points represent mean values with 95% confidence intervals.

**Fig. 3.**
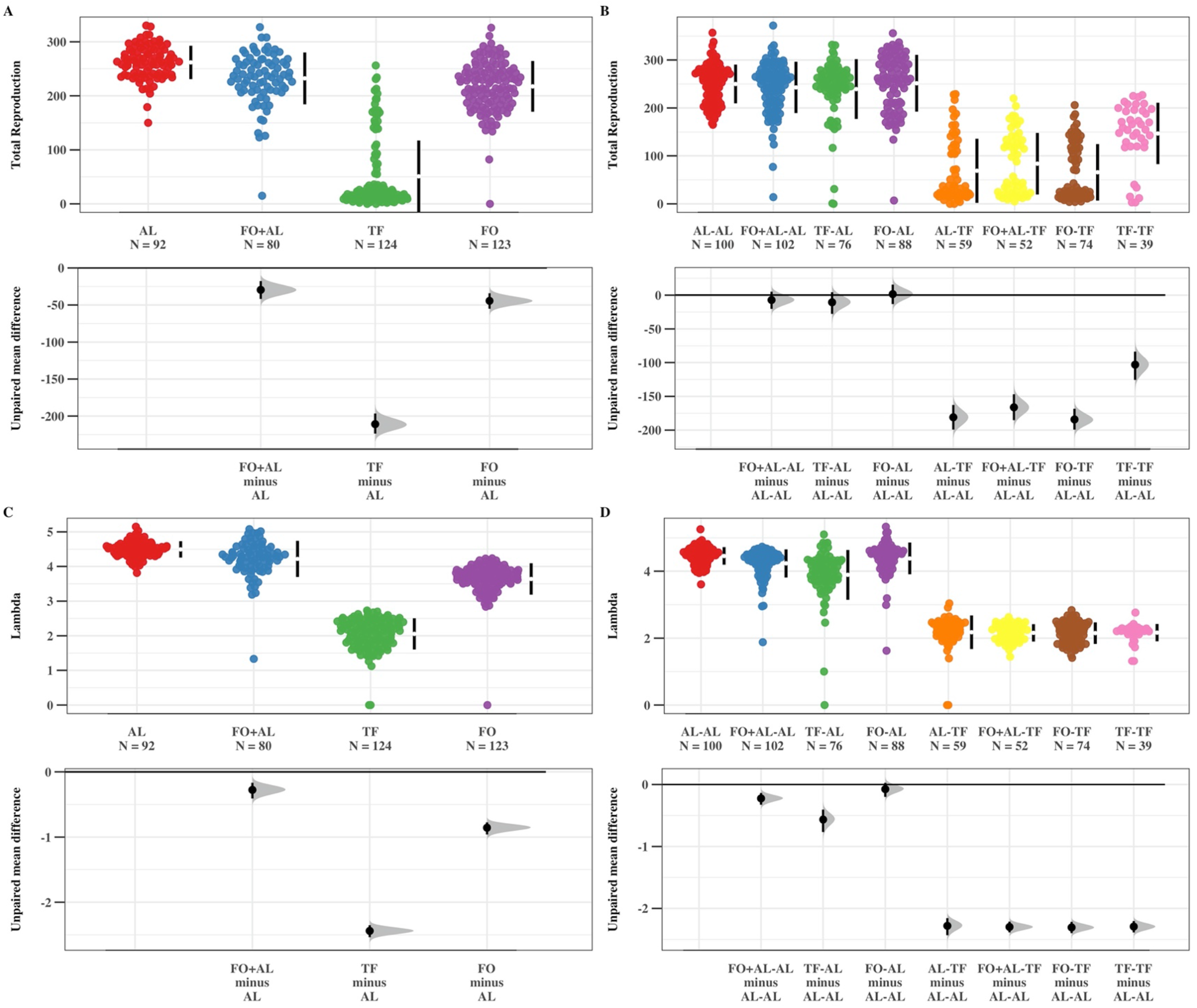
**A-D** Plots of LRS (A-B) and lambda (C-D) for P_0_ (A, C) and F_1_ (B, D). Colours represent differing dietary treatments for each generation. For Graphs B and D, the lineages are given as P_0_-F_1_. Bootstrapped mean comparison between treatments, showing mean values with 95% confidence intervals in both top and bottom panels.

**Fig. 4.**
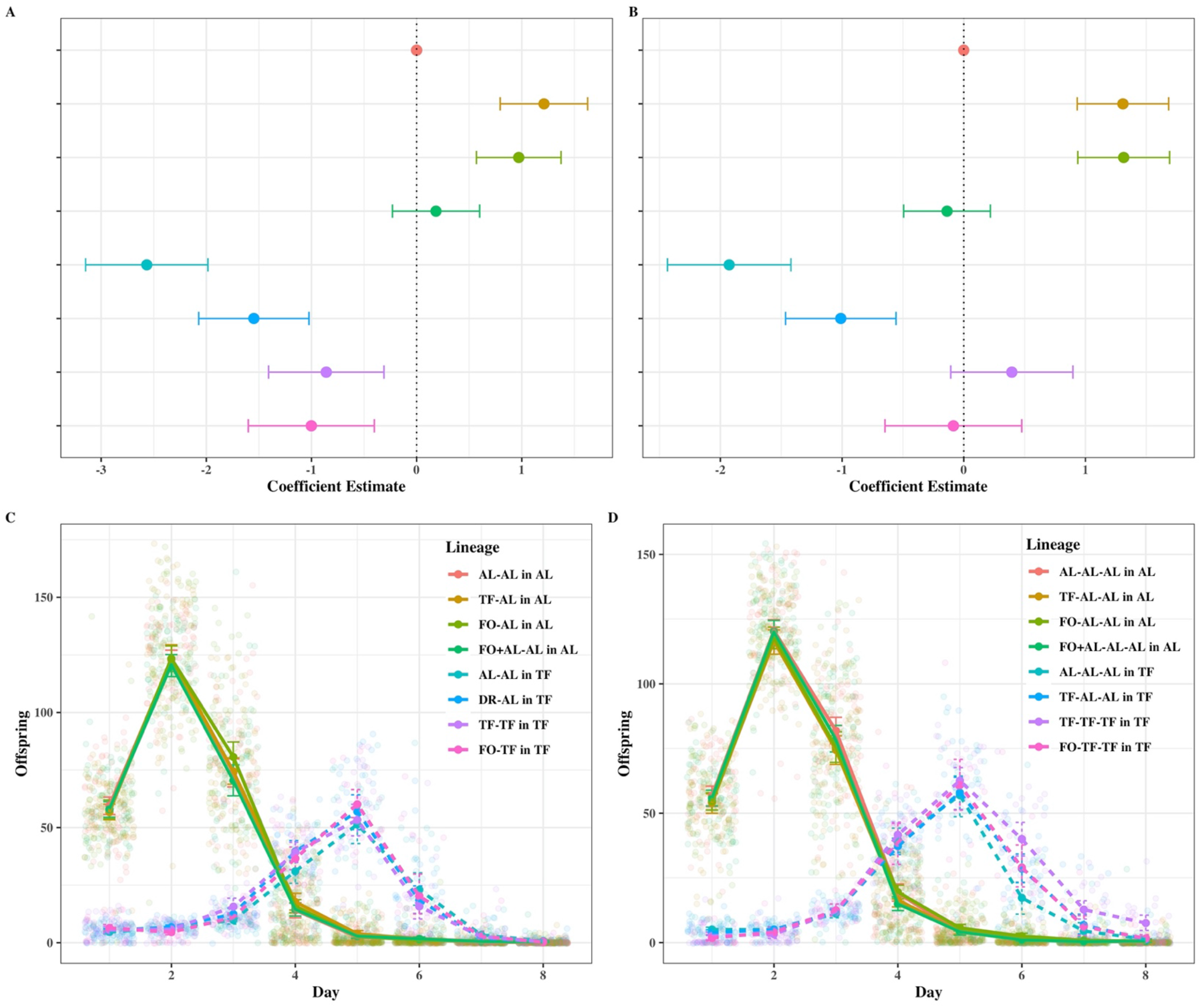
**A-D** Plots of survival (A-B) and reproduction (C-D) for F_2_ (A, C) and F_3_ (B, D). Colours represent differing dietary treatments for each generation. For graph C the lineages are given as P_0_-F_1_ in F_2_, whilst for graph D the lineages are given as P_0_-F_1_-F_2_ in F_3_. A-B) Points represent coefficients from a mixed effects cox model with 95% confidence intervals; C-D) Points represent mean values with 95% confidence intervals.

**Fig. 5.**
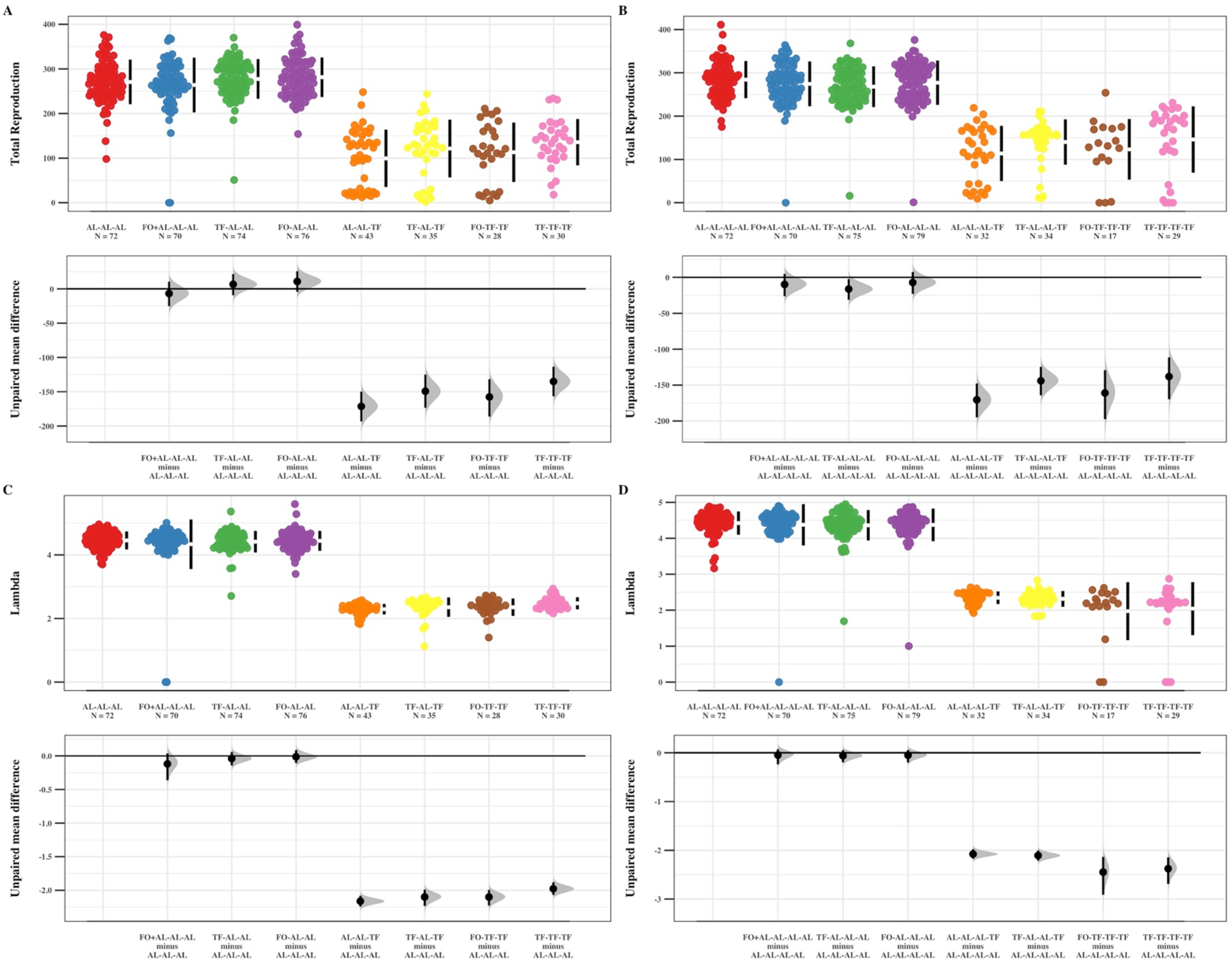
**A-D** Plots of LRS (A-B) and lambda (C-D) for F_2_ (A, C) and F_3_ (B, D). Colours represent differing dietary treatments for each generation. For graphs A and C, the lineages are given as P_0_-F_1_-F_2_, whilst for graphs B and D lineages are given as P_0_-F_1_-F_2_-F_3_. A-D) Bootstrapped comparison between treatments, showing mean values with 95% confidence intervals in both top and bottom panels.

Every generation was assayed for both lifespan and fitness by transferring onto new plates every 24 hours up until day eight and kept in climate chambers set to 20°C, 60%RH with continual darkness. After day eight, all surviving worms were then haphazardly positioned on top of a clear plastic tray and placed on a shelf at random within an incubator at 20°C with a 8h: 16g L:D cycle (the strength of light was ~7000 Lux) and transferred every two days onto new plates. Eggs laid within the eight-day reproductive period were allowed to hatch and develop for two days before hatched larvae were killed at 42°C for 3.5 hours and subsequently counted. For lifespan, death was defined as the absence of worm movement in response to touch. A worm was censored if the time of death was unknown, if it went missing, or if the worm gained access to the layer of food in the middle of the plate. These assays were repeated over three separate blocks. The experimental design of the first block differed slightly, with individual worms placed in light from Day 1 of reproduction, although the larvae still developed in darkness. In addition, this experimental block only continued until the F_1_ generation. As all treatments were otherwise treated similarly, with or without light from Day 1 of reproduction, we kept these Block 1 data within the analysis (See Figs. S1A-D and S2A-D).

### Statistical analysis

All analyses were performed using R v 3.6.0 (R Core Team, 2016).

Data used for the survival and reproduction analysed differed slightly. Whilst both datasets contained data from individuals where the time of death was known, individuals that died due to matricide were omitted from the survival analysis. For reproduction, these individuals were included but censored individuals (i.e. those where the exact time of death was not known) were omitted.

For all models involving P0, a generation-level factor of “Treatment” was added as a fixed effect with a random effect of “Founder” in order to account for possible pseudoreplication nested within each experimental “Block”. For models involving F_1_-F_3_ we fitted a new fixed covariate of dietary “Lineage” and a random effect of “Parent ID” (again, nested within “Block”).

Lifespan was analysed using Mixed Effect Cox Proportional Hazard Models from the coxme package v2.2-16 (Therneau, 2012) and visualised using Forest Plots for Hazard Ratios from the survminer package v0.4.6 (Kassambara, 2018).

For reproduction, three measures of fitness were analysed. Typically, age-specific reproduction within C*. elegans* nematodes is underdispersed with significant evidence of zero-inflation, particularly when individuals are dietary restricted. If these measures were indeed identified as zero-inflated (by simulating the residuals of a full Poisson model and testing for zero-inflation using the DHARMa package v 0.2.7, see Hartig 2020) an additional zero-inflated component and a variety of error distributions were fitted using the glmmTMB package v1.00 (Brooks *et al.*, 2017; Magnusson *et al.*, 2019). We fitted similar covariates as the survival models, however we added both the quadratic and linear fixed effects of “Day” and their interaction with either “Treatment” (for P_0_) or “Lineage” (for F_1_-F_3_) in order to capture the change in reproduction for each diet (or dietary lineage) over time. In addition, we also added a random effect of “Worm ID” in order to account for repeatedly measuring the same individual over multiple time points. Age-specific reproductive curves were visualised using ggplot2 (Wickham, 2009).

The second measure of fitness was total number of offspring produced by each individual, or Lifetime Reproductive Success (LRS). Similar fixed and random effects were used as in the mixed effects survival model. Mean differences between treatments were also analysed by bootstrapping and displayed in estimation plots from the dabestR package v0.2.4 (Ho *et al.*, 2019).

The last measure of fitness was individual fitness or *lind* and was obtained by constructing individual-based age-structured matrices (Leslie matrices, Leslie, 1945) and calculating the dominant eigenvalue using the lambda function from the popbio package v2.7 (Stubben & Milligan, 2007). Two days of development time was added onto the fertility schedule and represented time from egg to adulthood. These values were then analysed using glmmTMB with a Gaussian error structure and with similar factors as the previous models. Similarly, individual fitness was visualised on bootstrapped estimation plots from the dabestR package.

In all cases, aside from the individual fitness and survival model, model selection was performed to identify the best fitting error distribution and zero-inflation parameters for each response variable; chosen as the model with the lowest Akaike’s information criterion (AIC, see supplementary material for full model selection tables, S1A-T). In addition to the best fitting full model, partial models were also created in which data was subsetted into the two dietary conditions (TF and AL) in order to more clearly determine the effects of great-grand-/grand-/parental diets acting on offspring fitness.

## Results

### P_0_ Survival

The survival of P_0_ individuals was significantly impacted by their dietary treatment. Individuals that were temporarily fasted (TF) exhibited a significant reduction in mortality, and thus increased lifespan, in comparison to *ad libitum* (AL) individuals (TF: *β* = −1.104, 95%: −1.490, −0.717, *p*<0.001; Fig. 2A, Table S2A). In addition, whilst still significantly reduced in comparison to AL and FO+AL, individuals that were simply given the odour of food (FO) had mortality rates significantly greater than temporarily fasted individuals (FO: *β* = −0.356, 95%: −0.633, −0.079, *p* = 0.018; Fig. 2A, Table S2A).

### P_0_ Reproduction

In a similar manner to survival, all reproductive measures were significantly affected by dietary treatment (Table S2B-D). TF individuals exhibited a much delayed and reduced reproductive schedule in comparison to the FO treatment, which had an intermediate effect (Fig. 2C). In particular, these FO individuals exhibited a delayed reproductive peak in comparison to the FO+AL and AL treatments, but a far advanced peak when compared to TF individuals, who peaked on day five. Moreover, FO individuals produced a far greater total number of offspring in comparison to the TF treatment (FO: *β* = 1.99 (0.075), *p*<0.001; Fig. 3A, Table S2B-D). This earlier peak in reproduction also resulted in higher individual fitness for the FO treatment relative to TF individuals (FO: *β* = 1.54 (0.056), *p*<0.001; Fig. 3C, Table S2B-D).

### F_1_ Survival

Survival in the F_1_ generation was significantly affected by individual treatment, with TF individuals exhibiting reduced mortality risk in comparison to the AL treatment (Fig. 2B). Parental diet had no significant effect on individuals that were TF. However, in contrast, both the offspring from FO and FO+AL parents exhibited decreased mortality when placed in AL conditions (FO: *ß* = −0.691, 95%: −1.032, −0.351, *p*<0.001; FO+AL: *β* = −0.323, 95%: −0.644, −0.002, p = 0.048; Fig. 2B, Table S2E).

### F_1_ Reproduction

Parental diet significantly affected offspring reproduction and fitness in both the TF and AL environments (Table S2F-N, Fig. 2D, Fig. 3B,D). In AL conditions, both the LRS and individual fitness were reduced in offspring from TF parents in comparison to offspring from AL parents (*β* = −0.129 (0.036), *p* = <0.001; *ß* = - 0.660 (0.072), *p*<0.001; Fig. 3B,D, Table S2F-N). The opposite pattern was shown for individuals placed under TF. LRS was positively affected by parental treatment, with TF parents producing offspring with higher total reproductive count than offspring from AL parents (*β* = 0.448 (0.148), *p* = 0.003; Fig. 3B, Table S2F-N). However, as the reproductive peak was on day five rather than on days one and two, no detectable differences were found in individual fitness (*β* = −0.081 (0.078), *p* = 0.300, Fig. 3D, Table S2F-N). Offspring from FO parents exhibited similar detrimental effects when placed in mismatched environments as offspring from AL parents with reduced LRS in DR and no effect when placed in AL.

### F_2_ Survival

Survival was significantly impacted by grandparental dietary treatment. Grandparents that were placed within TF or FO conditions produced grand-offspring that exhibited a significant *increase* in mortality when placed within AL conditions (**TF**-AL-AL: *β* = 1.211, 95%: 0.795, 1.627, *p* <0.001; **FO**-AL-AL: *β* = 0.972, 95%: 0.569, 1.374, *p* <0.001; Fig. 4A, Table S2O) but a significant *decrease* in mortality when placed within TF conditions (**TF**-TF-TF: *β* = −0.859, 95%: −1.408, −0.310, *p* = 0.002; **FO**-TF-TF: *β* = −1.001, 95%: −1.600, - 0.402, *p* = 0.001; Fig. 4A, Table S2O). However, offspring from AL grand-parents and AL parents, with only one generation of fasting, exhibited increased lifespan in comparison to these two treatments (**AL**-AL-TF: *β* = −2.566, 95%: −3.148, −1.985, *p*<0.001; Fig. 4A, Table S2O).

### F_2_ Reproduction

Grandparental diet had no significant effect on any of the measured reproductive traits when individuals were placed in AL conditions (Table S2P-X). However, the mean values suggested that both TF and FO grandparents produced grand-offspring with increased average LRS (**TF**-AL-AL and **FO**-AL-AL). In the TF conditions, TF grandparents produced grand-offspring with significantly higher LRS in comparison to individuals produced from AL grandparents (**AL**-AL-TF: *β* = −0.3921 (0.1654), *p* = 0.0178; Fig. 5A, Table S2P-X). In addition, the full model revealed significantly lowered LRS in offspring produced from both FO and TF grandparents *with* AL parents (**FO**-TF-TF: *β* = −0.349 (0.113), *p* = 0.002; **TF**-AL-TF: *β* = −0.255 (0.104) *p* = 0.0142; Fig. 5A, Table S2P-X). Lastly, the partial model of individual fitness within the TF environment revealed a detectable decrease in lambda for offspring born from AL grandparents (**TF**-AL-AL: *β* = −0.1815 (0.061), *p* = 0.003; Fig. 5C, Table S2P-X).

### F_3_ Survival

In a similar manner to F_2_, survival of individuals was negatively impacted by great-grandparental treatment. With great-grandparents that were placed within TF or FO conditions producing great-grand-offspring with increased mortality risk in comparison to other treatments (**TF**-AL-AL-AL: *β* = 1.308, 95%: 0.933, 1.684, *p*<0.001; **FO**-AL-AL-AL: *β* = 1.314, 95%: 0.937, 1.692, *p*<0.001; Fig. 4B, Table S2Y). Interestingly, the positive effects of TF, which prevailed throughout the previous generations, disappeared. With the cumulative effects of four successive generations (three for the FO great-grandparental treatment) in TF environments causing a distinct lack of lifespan increase (**TF**-TF-TF-TF: *β* = 0.395, 95%: −0.107, 0.898, *p* = 0.123; **FO**-TF-TF-TF: *β* = −0.085, 95%: −0.647, 0.477, *p* = 0.766; Fig. 4B, Table S2Y).

### F_3_ Reproduction

Great-grandparental diet had detectable transgenerational effects acting on the LRS and fitness of individuals in both the AL and TF environments (Table S2Z-AH and Figs. 4D and 5B/D). Great-grand-offspring of TF individuals exhibited decreased LRS in AL environments compared to the AL control lineage (**AL**-AL-AL-AL, **TF**-AL-AL-AL: *β* = −0.060 (0.028), *p* = 0.032; Fig. 5B, Table S2Z-AH). In addition, great-grand-offspring of AL individuals exhibited decreased LRS in TF environments compared to the cumulative TF treatment (**TF**-TF-TF-TF, **AL**-AL-AL-TF: *β* = −0.352 (0.152), *p* = 0.021; Fig. 5B, Table S2Z-AH). For individual fitness, there were no detectable transgenerational effects when individuals were placed in AL environments, however individuals exhibited decreased lambda values when placed within TF environments. In particular, individuals produced from three or more generations of TF, exhibited significantly lowered fitness, including great-grand-offspring produced from four generations of TF and from FO great-grandparents (**TF**-TF-TF-TF: *β* = −0.318 (0.127), p = 0.012; **FO**-TF-TF-TF: *β* = −0.399 (0.151), *p* = 0.0084; Fig. 5D, Table S2Z-AH).

## Discussion

Mostly strikingly, we found evidence of several deleterious transgenerational effects of DR by temporary fasting in P_0_ on survival and fitness of individuals from the F_3_ generation. Specifically, great-grandparental DR increased mortality risk and reduced fitness of F_3_ descendants. Contrary to previous work showing that larval starvation can produce positive transgenerational effects on lifespan in *C. elegans*, we demonstrate that even temporary fasting in adults can result in detrimental transgenerational effects on lifespan by increasing mortality risk. Moreover, individuals produced from three generations of temporary fasting no longer displayed the classical reduction in mortality risk associated with DR and exhibited significantly reduced individual fitness. Taken together, these results highlight previously unknown long-term costs of DR which may have significant detrimental effects that only manifest in distant generations.

Beside the clear inter- and transgenerational trade-offs associated with the DR response, we also found that olfactory cues affect somatic maintenance and survival across several generations. P_0_ individuals given the odour of food but placed in the same environment as DR individuals, exhibited increased reproduction at the cost of reduced lifespan extension. Moreover, F_1_ offspring produced from food-odour DR parents behaved in a similar manner to offspring born from *ad libitum* parents. Not only does this suggest that parental effects regarding dietary condition are reliant upon accurate environmental cues being passed to the next generation, but also that investment into survival is largely in response to food-related odours which promote longevity through the activation of insulin/IGF-1 pathways (Alcedo & Kenyon, 2004). It is possible that nematodes are reluctant to lay eggs in the environment perceived as devoid of food and that reduced reproduction under DR is driven partly by the lack of resources to produce gametes and also as a result of an adaptive reproductive strategy by the organism. Taken together, these results are largely in line with the resource reallocation hypothesis (Shanley & Kirkwood, 2000) where individuals under DR reallocate resources from reproduction into somatic maintenance to increase the probability of survival until the next reproductive opportunity.

Our results clearly answer a number of important questions regarding lifespan extension via dietary restriction in parents. Consistent with previous research (Houthoofd *et al.* 2007; Smith *et al.* 2008; Mautz *et al.* 2020, and see Nakagawa *et al.* 2012), DR via temporary fasting resulted in a detectable reduction in mortality risk at the cost of reduced reproduction in the P_0_ generation. Moreover, we found when individuals were placed back onto a standard food regime post-DR (Day 3), reproduction steadily increased until a peak occurred on Day 5. This contrasts with the findings of McCracken *et al.* (2020) who found that the survival and fertility of *Drosophila melanogaster* decreased immediately following a return to a rich diet after a period of DR, however they note that integral to this increase is the relative duration of DR prior to rich feeding. It is possible therefore, that we may have seen similar patterns of mortality exacerbation if nematodes remained within this temporary fasting environment for longer.

Our results also showed that DR in P_0_ not only affects the present generation but also has long-term effects for up to three subsequent generations. We identified significant long-term effects of both parental and ancestral diet manifesting in changes to survival and fitness of subsequent generations. In fact, if the parental generation and F_1_ offspring were exposed to the same dietary treatment, the benefits carried on across both generations. This finding suggests the transmission of information allows offspring to anticipate potentially adverse conditions, such as dietary restriction, resulting in increased performance. Although different in methodology, these results are largely consistent with intergenerational phenotypic plasticity acting between mother and offspring identified by Hibshman *et al.* (2016). However, we extend this work by showing that offspring of DR parents pay a price of reduced fitness when raised in standard environmental conditions. Thus, the parental DR response can be costly for the offspring and its adaptive nature depends on whether the current environment of the parents matches the future environment of their offspring.

## Concluding Remarks

Dietary restriction (DR) by temporary fasting produced a wide variety of both beneficial and deleterious effects across four generations. Whilst the effects of DR were positive for survival in the P_0_ generation, and for both fitness and survival in the F_1_ generation in matching environments, we revealed deleterious transgenerational effects on survival and reproduction in both the F_2_ and F_3_ generations. Strikingly, our results also reveal negative transgenerational effects on survival and fitness as a result of accumulated DR exposure across multiple generations. Furthermore, the beneficial effects of DR acting to increase lifespan and reduce mortality risk, all but vanish when considering F_3_ individuals. Taken together, these results highlight important long-term costs of lifespan extension via DR. Future research should focus on discovering DR regimes that robustly extend healthy lifespan while avoiding negative consequences for future generations of descendants.

## Authors’ Contributions

EIC and AAM conceived the project and EIC, KS, HC, SI, TC, and AAM designed the experiments and the methodology. EIC analysed the data. EIC and AAM wrote the manuscript; KS, HC, SI and TC commented on the drafts. EIC and KS collected the data. All authors gave final approval for publication.

## Acknowledgements

This work was funded by BBSRC BB/R017387/1 and ERC GermlineAgeingSoma 724909 to AAM.

Thanks go to members of the Maklakov Lab and Eryn Macfarlane for statistical advice. The authors declare no competing interests.

