## Supplementary for "Transgenerational fitness effects of lifespan extension by dietary restriction in *Caenorhabditis elegans*"

**Table S1A.** Model Selection for Age-Specific Reproduction in $P_0$.

*N.B.* The model in bold denotes the best fitting model with the lowest AIC from the glmmTMB package. Models with NAs failed to converge.

**Significant zero-inflation was identified in a standard Poisson model (DHARMa zero-inflation test *p* = <0.001*)***

| Model | Family | Model formula | Zero-inflation formula | df | Loglik | AIC | Delta AIC |
|---|---|---|---|---|---|---|---|
| **m32** | **genpois(log)** | **~Treatment*Day2+ Treatment*Day + (1\|Block/Founder/Worm)** | **~Treatment*Day2+Treatment*Day** | **28** | **-9329.7722** | **18715.5444** | **0** |
| m30 | genpois(log) | ~Treatment*Day2+ Treatment*Day + (1\|Block/Founder/Worm) | ~Treatment*Day+Day2 | 25 | -9336.7764 | 18723.5528 | 8.0084682 |
| m31 | genpois(log) | ~Treatment*Day2+ Treatment*Day + (1\|Block/Founder/Worm) | ~Treatment*Day2+Day | 25 | -9337.2105 | 18724.421 | 8.87668896 |
| m12 | nbinom1(log) | ~Treatment*Day2+ Treatment*Day + (1\|Block/Founder/Worm) | ~Treatment+Day | 21 | -9351.9214 | 18745.8428 | 30.2984072 |
| m13 | nbinom1(log) | ~Treatment*Day2+ Treatment*Day + (1\|Block/Founder/Worm) | ~Treatment+Day+Day2 | 22 | -9351.8305 | 18747.661 | 32.1166885 |
| m28 | genpois(log) | ~Treatment*Day2+ Treatment*Day + (1\|Block/Founder/Worm) | ~Treatment+Day | 21 | -9358.1506 | 18758.3012 | 42.7568882 |
| m29 | genpois(log) | ~Treatment*Day2+ Treatment*Day + (1\|Block/Founder/Worm) | ~Treatment+Day+Day2 | 22 | -9357.9724 | 18759.9448 | 44.4004686 |
| m10 | nbinom1(log) | ~Treatment*Day2+ Treatment*Day + (1\|Block/Founder/Worm) | ~1 | 17 | -9373.9066 | 18781.8132 | 66.2688696 |
| m11 | nbinom1(log) | ~Treatment*Day2+ Treatment*Day + (1\|Block/Founder/Worm) | ~Treatment | 20 | -9371.6858 | 18783.3716 | 67.8272154 |
| m26 | genpois(log) | ~Treatment*Day2+ Treatment*Day + (1\|Block/Founder/Worm) | ~1 | 17 | -9382.8959 | 18799.7918 | 84.2474835 |
| m27 | genpois(log) | ~Treatment*Day2+ Treatment*Day + (1\|Block/Founder/Worm) | ~Treatment | 20 | -9380.9308 | 18801.8615 | 86.3171532 |
| m9 | nbinom1(log) | ~Treatment*Day2+ Treatment*Day + (1\|Block/Founder/Worm) | ~0 | 16 | -9427.344 | 18886.688 | 171.143678 |
| m25 | genpois(log) | ~Treatment*Day2+ Treatment*Day + (1\|Block/Founder/Worm) | ~0 | 16 | -9444.7117 | 18921.4234 | 205.879025 |
| m24 | nbinom2(log) | ~Treatment*Day2+ Treatment*Day + (1\|Block/Founder/Worm) | ~Treatment*Day2+Treatment*Day | 28 | -9704.0301 | 19464.0601 | 748.515751 |
| m20 | nbinom2(log) | ~Treatment*Day2+ Treatment*Day + (1\|Block/Founder/Worm) | ~Treatment+Day | 21 | -9752.4739 | 19546.9477 | 831.403357 |
| m19 | nbinom2(log) | ~Treatment*Day2+ Treatment*Day + (1\|Block/Founder/Worm) | ~Treatment | 20 | -9937.9759 | 19915.9517 | 1200.40738 |
| m18 | nbinom2(log) | ~Treatment*Day2+ Treatment*Day + (1\|Block/Founder/Worm) | ~1 | 17 | -9942.3069 | 19918.6138 | 1203.06949 |
| m17 | nbinom2(log) | ~Treatment*Day2+ Treatment*Day + (1\|Block/Founder/Worm) | ~0 | 16 | -9975.9344 | 19983.8688 | 1268.32444 |
| m7 | poisson(log) | ~Treatment*Day2+ Treatment*Day + (1\|Block/Founder/Worm) | ~Treatment*Day2+Day | 24 | -14242.952 | 28533.9046 | 9818.36028 |
| m6 | poisson(log) | ~Treatment*Day2+ Treatment*Day + (1\|Block/Founder/Worm) | ~Treatment*Day+Day2 | 24 | -14245.238 | 28538.4759 | 9822.93157 |
| m5 | poisson(log) | ~Treatment*Day2+ Treatment*Day + (1\|Block/Founder/Worm) | ~Treatment+Day+Day2 | 21 | -14273.285 | 28588.569 | 9873.02468 |
| m4 | poisson(log) | ~Treatment*Day2+ Treatment*Day + (1\|Block/Founder/Worm) | ~Treatment+Day | 20 | -14280.47 | 28600.9409 | 9885.39659 |
| m3 | poisson(log) | ~Treatment*Day2+ Treatment*Day + (1\|Block/Founder/Worm) | ~Treatment | 19 | -14434.938 | 28907.8769 | 10192.3326 |
| m2 | poisson(log) | ~Treatment*Day2+ Treatment*Day + (1\|Block/Founder/Worm) | ~1 | 16 | -14450.139 | 28932.2787 | 10216.7343 |
| m1 | poisson(log) | ~Treatment*Day2+ Treatment*Day + (1\|Block/Founder/Worm) | ~0 | 15 | -15082.643 | 30195.2857 | 11479.7413 |

| | | | | | | | |
|---|---|---|---|---|---|---|---|
| m8 | poisson(log) | ~Treatment*Day2+ Treatment*Day + (1\|Block/Founder/Worm) | ~Treatment*Day2+Treatment*Day | 27 | NA | NA | NA |
| m14 | nbinom1(log) | ~Treatment*Day2+ Treatment*Day + (1\|Block/Founder/Worm) | ~Treatment*Day+Day2 | 25 | NA | NA | NA |
| m15 | nbinom1(log) | ~Treatment*Day2+ Treatment*Day + (1\|Block/Founder/Worm) | ~Treatment*Day2+Day | 25 | NA | NA | NA |
| m16 | nbinom1(log) | ~Treatment*Day2+ Treatment*Day + (1\|Block/Founder/Worm) | ~Treatment*Day2+Treatment*Day | 28 | NA | NA | NA |
| m21 | nbinom2(log) | ~Treatment*Day2+ Treatment*Day + (1\|Block/Founder/Worm) | ~Treatment+Day+Day2 | 22 | NA | NA | NA |
| m22 | nbinom2(log) | ~Treatment*Day2+ Treatment*Day + (1\|Block/Founder/Worm) | ~Treatment*Day+Day2 | 25 | NA | NA | NA |
| m23 | nbinom2(log) | ~Treatment*Day2+ Treatment*Day + (1\|Block/Founder/Worm) | ~Treatment*Day2+Day | 25 | NA | NA | NA |

**Table S1B.** Model Selection for Total Reproduction in $P_0$.

**Significant zero-inflation was identified in a standard Poisson model (DHARMa zero-inflation test *p* = <0.001*).***

| Model | Family | Model formula | Zero-inflation formula | df | Loglik | AIC | Delta AIC |
|---|---|---|---|---|---|---|---|
| **m6.1** | **genpois(log)** | **~Treatment + (1\|Block/Founder)** | **~1** | **8** | **-2234.0753** | **4484.15061** | **0** |
| m7 | genpois(log) | ~Treatment + (1\|Block/Founder) | ~Treatment | 11 | -2232.739 | 4487.47802 | 3.32740859 |
| m2.1 | nbinom1(log) | ~Treatment + (1\|Block/Founder) | ~1 | 8 | -2240.2222 | 4496.44444 | 12.2938278 |
| m6 | genpois(log) | ~Treatment + (1\|Block/Founder) | ~0 | 7 | -2276.2386 | 4566.47716 | 82.3265452 |
| m4.1 | nbinom2(log) | ~Treatment + (1\|Block/Founder) | ~1 | 8 | -2416.6889 | 4849.37777 | 365.227156 |
| m5 | nbinom2(log) | ~Treatment + (1\|Block/Founder) | ~Treatment | 11 | -2415.036 | 4852.0721 | 367.921483 |
| m4 | nbinom2(log) | ~Treatment + (1\|Block/Founder) | ~0 | 7 | -2425.9017 | 4865.80335 | 381.652734 |
| m1.2 | poisson(log) | ~Treatment + (1\|Block/Founder) | ~1 | 7 | -6234.4829 | 12482.9658 | 7998.81521 |
| m1.3 | poisson(log) | ~Treatment + (1\|Block/Founder) | ~Treatment | 10 | -6232.7221 | 12485.4441 | 8001.29351 |
| m1.1 | poisson(log) | ~Treatment + (1\|Block/Founder) | ~0 | 6 | -6539.645 | 13091.2901 | 8607.13946 |
| m2 | nbinom1(log) | ~Treatment + (1\|Block/Founder) | ~0 | 7 | NA | NA | NA |
| m3 | nbinom1(log) | ~Treatment + (1\|Block/Founder) | ~Treatment | 11 | NA | NA | NA |

**Table S1C.** Full model selection for Age-Specific Reproduction in $F_1$.

**Significant zero-inflation was identified in a standard Poisson model (DHARMa zero-inflation test *p* = 0.016*).***

| Model | Family | Model formula | Zero-inflation formula | df | Loglik | AIC | Delta AIC |
|---|---|---|---|---|---|---|---|
| **m29** | **genpois(log)** | **~Lineage*Day2+ Lineage*Day + (1\|Block/Parent.No/Worm)** | **~Lineage+Day+Day2** | **38** | **-14316.096** | **28708.1919** | **0** |
| m12 | nbinom1(log) | ~Lineage*Day2+ Lineage*Day + (1\|Block/Parent.No/Worm) | ~Lineage+Day | 37 | -14325.843 | 28725.686 | 17.49402 |
| m11 | nbinom1(log) | ~Lineage*Day2+ Lineage*Day + (1\|Block/Parent.No/Worm) | ~Lineage | 36 | -14333.389 | 28738.7774 | 30.58544 |
| m28 | genpois(log) | ~Lineage*Day2+ Lineage*Day + (1\|Block/Parent.No/Worm) | ~Lineage+Day | 37 | -14334.891 | 28743.783 | 35.59103 |
| m10 | nbinom1(log) | ~Lineage*Day2+ Lineage*Day + (1\|Block/Parent.No/Worm) | ~1 | 29 | -14350.971 | 28759.942 | 51.75006 |
| m27 | genpois(log) | ~Lineage*Day2+ Lineage*Day + (1\|Block/Parent.No/Worm) | ~Lineage | 36 | -14347.461 | 28766.922 | 58.73003 |
| m26 | genpois(log) | ~Lineage*Day2+ Lineage*Day + (1\|Block/Parent.No/Worm) | ~1 | 29 | -14363.15 | 28784.3004 | 76.10841 |
| m9 | nbinom1(log) | ~Lineage*Day2+ Lineage*Day + (1\|Block/Parent.No/Worm) | ~0 | 28 | -14440.869 | 28937.7377 | 229.54576 |
| m25 | genpois(log) | ~Lineage*Day2+ Lineage*Day + (1\|Block/Parent.No/Worm) | ~0 | 28 | -14472.448 | 29000.8968 | 292.70487 |

| | | | | | | | |
|---|---|---|---|---|---|---|---|
| m21 | nbinom2(log) | ~Lineage*Day2+ Lineage*Day + (1\|Block/Parent.No/Worm) | ~Lineage+Day+Day2 | 38 | -14594.175 | 29264.3507 | 556.15871 |
| m20 | nbinom2(log) | ~Lineage*Day2+ Lineage*Day + (1\|Block/Parent.No/Worm) | ~Lineage+Day | 37 | -14645.532 | 29365.0648 | 656.87286 |
| m19 | nbinom2(log) | ~Lineage*Day2+ Lineage*Day + (1\|Block/Parent.No/Worm) | ~Lineage | 36 | -14888.585 | 29849.1691 | 1140.97713 |
| m18 | nbinom2(log) | ~Lineage*Day2+ Lineage*Day + (1\|Block/Parent.No/Worm) | ~1 | 29 | -14907.584 | 29873.1674 | 1164.97548 |
| m17 | nbinom2(log) | ~Lineage*Day2+ Lineage*Day + (1\|Block/Parent.No/Worm) | ~0 | 28 | -14934.146 | 29924.2922 | 1216.10027 |
| m6 | poisson(log) | ~Lineage*Day2+ Lineage*Day + (1\|Block/Parent.No/Worm) | ~Lineage*Day+Day2 | 44 | -23540.202 | 47168.4037 | 18460.2117 |
| m5 | poisson(log) | ~Lineage*Day2+ Lineage*Day + (1\|Block/Parent.No/Worm) | ~Lineage+Day+Day2 | 37 | -23601.345 | 47276.6901 | 18568.4982 |
| m4 | poisson(log) | ~Lineage*Day2+ Lineage*Day + (1\|Block/Parent.No/Worm) | ~Lineage+Day | 36 | -23608.079 | 47288.1588 | 18579.9668 |
| m7 | poisson(log) | ~Lineage*Day2+ Lineage*Day + (1\|Block/Parent.No/Worm) | ~Lineage*Day2+Day | 44 | -23606.868 | 47301.7367 | 18593.5448 |
| m3 | poisson(log) | ~Lineage*Day2+ Lineage*Day + (1\|Block/Parent.No/Worm) | ~Lineage | 35 | -23740.423 | 47550.8467 | 18842.6548 |
| m2 | poisson(log) | ~Lineage*Day2+ Lineage*Day + (1\|Block/Parent.No/Worm) | ~1 | 28 | -23754.097 | 47564.195 | 18856.003 |
| m1 | poisson(log) | ~Lineage*Day2+ Lineage*Day + (1\|Block/Parent.No/Worm) | ~0 | 27 | -25378.023 | 50810.0465 | 22101.8546 |
| m8 | poisson(log) | ~Lineage*Day2+ Lineage*Day + (1\|Block/Parent.No/Worm) | ~Lineage*Day2+Lineage*Day | 51 | NA | NA | NA |
| m13 | nbinom1(log) | ~Lineage*Day2+ Lineage*Day + (1\|Block/Parent.No/Worm) | ~Lineage+Day+Day2 | 38 | NA | NA | NA |
| m14 | nbinom1(log) | ~Lineage*Day2+ Lineage*Day + (1\|Block/Parent.No/Worm) | ~Lineage*Day+Day2 | 45 | NA | NA | NA |
| m15 | nbinom1(log) | ~Lineage*Day2+ Lineage*Day + (1\|Block/Parent.No/Worm) | ~Lineage*Day2+Day | 45 | NA | NA | NA |
| m16 | nbinom1(log) | ~Lineage*Day2+ Lineage*Day + (1\|Block/Parent.No/Worm) | ~Lineage*Day2+Lineage*Day | 52 | NA | NA | NA |
| m22 | nbinom2(log) | ~Lineage*Day2+ Lineage*Day + (1\|Block/Parent.No/Worm) | ~Lineage*Day+Day2 | 45 | NA | NA | NA |
| m23 | nbinom2(log) | ~Lineage*Day2+ Lineage*Day + (1\|Block/Parent.No/Worm) | ~Lineage*Day2+Day | 45 | NA | NA | NA |
| m24 | nbinom2(log) | ~Lineage*Day2+ Lineage*Day + (1\|Block/Parent.No/Worm) | ~Lineage*Day2+Lineage*Day | 52 | NA | NA | NA |
| m30 | genpois(log) | ~Lineage*Day2+ Lineage*Day + (1\|Block/Parent.No/Worm) | ~Lineage*Day+Day2 | 45 | NA | NA | NA |
| m31 | genpois(log) | ~Lineage*Day2+ Lineage*Day + (1\|Block/Parent.No/Worm) | ~Lineage*Day2+Day | 45 | NA | NA | NA |
| m32 | genpois(log) | ~Lineage*Day2+ Lineage*Day + (1\|Block/Parent.No/Worm) | ~Lineage*Day2+Lineage*Day | 52 | NA | NA | NA |

**Table S1D.** Partial model selection for Age-Specific Reproduction in $F_1$ in temporary fasting conditions.

**Significant zero-inflation was identified in a standard Poisson model (DHARMa zero-inflation test *p* = 0.048*).***

| Model | Family | Model formula | Zero-inflation formula | df | Loglik | AIC | Delta AIC |
|---|---|---|---|---|---|---|---|
| **m29** | **genpois(log)** | **~Lineage*Day2+ Lineage*Day + (1\|Block/Parent.No/Worm)** | **~Lineage+Day+Day2** | **22** | **-4711.3508** | **9466.70155** | **0** |
| m28 | genpois(log) | ~Lineage*Day2+ Lineage*Day + (1\|Block/Parent.No/Worm) | ~Lineage+Day | 21 | -4720.952 | 9483.90399 | 17.2024468 |
| m13 | nbinom1(log) | ~Lineage*Day2+ Lineage*Day + (1\|Block/Parent.No/Worm) | ~Lineage+Day+Day2 | 22 | -4726.897 | 9497.79394 | 31.0923973 |
| m14 | nbinom1(log) | ~Lineage*Day2+ Lineage*Day + (1\|Block/Parent.No/Worm) | ~Lineage*Day+Day2 | 25 | -4725.901 | 9501.80198 | 35.1004311 |
| m26 | genpois(log) | ~Lineage*Day2+ Lineage*Day + (1\|Block/Parent.No/Worm) | ~1 | 17 | -4735.8768 | 9505.75353 | 39.0519833 |
| m27 | genpois(log) | ~Lineage*Day2+ Lineage*Day + (1\|Block/Parent.No/Worm) | ~Lineage | 20 | -4733.5379 | 9507.07588 | 40.374335 |
| m25 | genpois(log) | ~Lineage*Day2+ Lineage*Day + (1\|Block/Parent.No/Worm) | ~0 | 16 | -4741.2907 | 9514.58134 | 47.8797919 |
| m12 | nbinom1(log) | ~Lineage*Day2+ Lineage*Day + (1\|Block/Parent.No/Worm) | ~Lineage+Day | 21 | -4737.341 | 9516.68204 | 49.9804891 |
| m10 | nbinom1(log) | ~Lineage*Day2+ Lineage*Day + (1\|Block/Parent.No/Worm) | ~1 | 17 | -4751.4154 | 9536.83077 | 70.1292242 |
| m9 | nbinom1(log) | ~Lineage*Day2+ Lineage*Day + (1\|Block/Parent.No/Worm) | ~0 | 16 | -4753.1324 | 9538.26479 | 71.5632454 |
| m11 | nbinom1(log) | ~Lineage*Day2+ Lineage*Day + (1\|Block/Parent.No/Worm) | ~Lineage | 20 | -4749.3397 | 9538.67946 | 71.9779169 |
| m21 | nbinom2(log) | ~Lineage*Day2+ Lineage*Day + (1\|Block/Parent.No/Worm) | ~Lineage+Day+Day2 | 22 | -4755.7192 | 9555.4384 | 88.7368532 |
| m20 | nbinom2(log) | ~Lineage*Day2+ Lineage*Day + (1\|Block/Parent.No/Worm) | ~Lineage+Day | 21 | -4759.3643 | 9560.72852 | 94.02697 |

| | | | | | | | |
|---|---|---|---|---|---|---|---|
| m19 | nbinom2(log) | ~Lineage*Day2+ Lineage*Day + (1\|Block/Parent.No/Worm) | ~Lineage | 20 | -4767.8166 | 9575.6333 | 108.931751 |
| m18 | nbinom2(log) | ~Lineage*Day2+ Lineage*Day + (1\|Block/Parent.No/Worm) | ~1 | 17 | -4771.2745 | 9576.54903 | 109.847483 |
| m17 | nbinom2(log) | ~Lineage*Day2+ Lineage*Day + (1\|Block/Parent.No/Worm) | ~0 | 16 | -4773.8374 | 9579.67486 | 112.973312 |
| m6 | poisson(log) | ~Lineage*Day2+ Lineage*Day + (1\|Block/Parent.No/Worm) | ~Lineage*Day+Day2 | 24 | -6750.7073 | 13549.4146 | 4082.71302 |
| m7 | poisson(log) | ~Lineage*Day2+ Lineage*Day + (1\|Block/Parent.No/Worm) | ~Lineage*Day2+Day | 24 | -6753.593 | 13555.186 | 4088.48444 |
| m5 | poisson(log) | ~Lineage*Day2+ Lineage*Day + (1\|Block/Parent.No/Worm) | ~Lineage+Day+Day2 | 21 | -6760.7194 | 13563.4388 | 4096.73729 |
| m4 | poisson(log) | ~Lineage*Day2+ Lineage*Day + (1\|Block/Parent.No/Worm) | ~Lineage+Day | 20 | -6776.5973 | 13593.1945 | 4126.49298 |
| m8 | poisson(log) | ~Lineage*Day2+ Lineage*Day + (1\|Block/Parent.No/Worm) | ~Lineage*Day2+Lineage*Day | 27 | -6847.751 | 13749.5021 | 4282.80055 |
| m2 | poisson(log) | ~Lineage*Day2+ Lineage*Day + (1\|Block/Parent.No/Worm) | ~1 | 16 | -6872.1219 | 13776.2437 | 4309.54217 |
| m3 | poisson(log) | ~Lineage*Day2+ Lineage*Day + (1\|Block/Parent.No/Worm) | ~Lineage | 19 | -6871.2347 | 13780.4693 | 4313.76776 |
| m1 | poisson(log) | ~Lineage*Day2+ Lineage*Day + (1\|Block/Parent.No/Worm) | ~0 | 15 | -7274.8326 | 14579.6651 | 5112.96358 |
| m15 | nbinom1(log) | ~Lineage*Day2+ Lineage*Day + (1\|Block/Parent.No/Worm) | ~Lineage*Day2+Day | 25 | NA | NA | NA |
| m16 | nbinom1(log) | ~Lineage*Day2+ Lineage*Day + (1\|Block/Parent.No/Worm) | ~Lineage*Day2+Lineage*Day | 28 | NA | NA | NA |
| m22 | nbinom2(log) | ~Lineage*Day2+ Lineage*Day + (1\|Block/Parent.No/Worm) | ~Lineage*Day+Day2 | 25 | NA | NA | NA |
| m23 | nbinom2(log) | ~Lineage*Day2+ Lineage*Day + (1\|Block/Parent.No/Worm) | ~Lineage*Day2+Day | 25 | NA | NA | NA |
| m24 | nbinom2(log) | ~Lineage*Day2+ Lineage*Day + (1\|Block/Parent.No/Worm) | ~Lineage*Day2+Lineage*Day | 28 | NA | NA | NA |
| m30 | genpois(log) | ~Lineage*Day2+ Lineage*Day + (1\|Block/Parent.No/Worm) | ~Lineage*Day+Day2 | 25 | NA | NA | NA |
| m31 | genpois(log) | ~Lineage*Day2+ Lineage*Day + (1\|Block/Parent.No/Worm) | ~Lineage*Day2+Day | 25 | NA | NA | NA |
| m32 | genpois(log) | ~Lineage*Day2+ Lineage*Day + (1\|Block/Parent.No/Worm) | ~Lineage*Day2+Lineage*Day | 28 | NA | NA | NA |

**Table S1E.** Partial model selection for Age-Specific Reproduction in $F_1$ in *ad libitum* conditions.

**No significant zero-inflation was identified in a standard Poisson model (DHARMa zero-inflation test *p* = 0.192*).***

| Model | Family | Model formula | df | Loglik | AIC | Delta AIC |
|---|---|---|---|---|---|---|
| **m9** | **nbinom1(log)** | **~Lineage*Day2+ Lineage*Day + (1\|Block/Parent.No/Worm)** | **16** | **-9601.8465** | **19235.6931** | **0** |
| m25 | genpois(log) | ~Lineage*Day2+ Lineage*Day + (1\|Block/Parent.No/Worm) | 16 | -9672.8308 | 19377.6615 | 141.968416 |
| m17 | nbinom2(log) | ~Lineage*Day2+ Lineage*Day + (1\|Block/Parent.No/Worm) | 16 | -10123.42 | 20278.8408 | 1043.14773 |
| m1 | poisson(log) | ~Lineage*Day2+ Lineage*Day + (1\|Block/Parent.No/Worm) | 15 | -17968.985 | 35967.9709 | 16732.2778 |

**Table S1F.** Full model selection for Total Reproduction in $F_1$.

**Significant zero-inflation was identified in a standard Poisson model (DHARMa zero-inflation test *p* = <0.001*).***

| Model | Family | Model formula | Zero-inflation formula | df | Loglik | AIC | Delta AIC |
|---|---|---|---|---|---|---|---|
| **m5** | **nbinom1(log)** | **~Lineage + (1\|BlockParent.No)** | **~1** | **12** | **-3248.6734** | **6521.34678** | **0** |
| m6 | nbinom1(log) | ~Lineage + (1\|BlockParent.No) | ~Lineage | 19 | -3243.9221 | 6525.84413 | 4.49735855 |
| m4 | nbinom1(log) | ~Lineage + (1\|BlockParent.No) | ~0 | 11 | -3270.0213 | 6562.04261 | 40.6958312 |
| m11 | genpois(log) | ~Lineage + (1\|BlockParent.No) | ~1 | 12 | -3295.8065 | 6615.61293 | 94.2661581 |
| m12 | genpois(log) | ~Lineage + (1\|BlockParent.No) | ~Lineage | 19 | -3291.0313 | 6620.06252 | 98.7157403 |

| | | | | | | | |
|---|---|---|---|---|---|---|---|
| m10 | genpois(log) | ~Lineage + (1\|BlockParent.No) | ~0 | 11 | -3318.8821 | 6659.76428 | 138.417499 |
| m8 | nbinom2(log) | ~Lineage + (1\|BlockParent.No) | ~1 | 12 | -3368.8783 | 6761.75669 | 240.409912 |
| m9 | nbinom2(log) | ~Lineage + (1\|BlockParent.No) | ~Lineage | 19 | -3364.1011 | 6766.20212 | 244.855345 |
| m7 | nbinom2(log) | ~Lineage + (1\|BlockParent.No) | ~0 | 11 | -3390.8683 | 6803.73652 | 282.389744 |
| m2 | poisson(log) | ~Lineage + (1\|BlockParent.No) | ~1 | 11 | -6157.8273 | 12337.6546 | 5816.30785 |
| m3 | poisson(log) | ~Lineage + (1\|BlockParent.No) | ~Lineage | 18 | -6153.049 | 12342.0981 | 5820.75131 |

**Table S1G.** Partial model selection for Total Reproduction in $F_1$ in temporary fasting conditions.

**No significant zero-inflation was identified in a standard Poisson model (DHARMa zero-inflation test *p* = 0.056*).***

| Model | Family | Model formula | df | Loglik | AIC | Delta AIC |
|---|---|---|---|---|---|---|
| **m7** | **nbinom2(log)** | **~Lineage + (1\|BlockParent.No)** | **7** | **-1146.6505** | **2307.3009** | **0** |
| m4 | nbinom1(log) | ~Lineage + (1\|BlockParent.No) | 7 | -1163.1547 | 2340.30949 | 33.0085857 |
| m10 | genpois(log) | ~Lineage + (1\|BlockParent.No) | 7 | -1168.7867 | 2351.57345 | 44.2725515 |
| m0 | poisson(log) | ~Lineage + (1\|BlockParent.No) | 6 | -1248.8861 | 2509.77215 | 202.471249 |

**Table S1H.** Partial model selection for Total Reproduction in $F_1$ in *ad libitum* conditions.

**Significant zero-inflation was identified in a standard Poisson model (DHARMa zero-inflation test *p* = <0.001*).***

| Model | Family | Model formula | Zero-inflation formula | df | Loglik | AIC | Delta AIC |
|---|---|---|---|---|---|---|---|
| **m5** | **nbinom1(log)** | **~Lineage + (1\|BlockParent.No)** | **~1** | **8** | **-1993.1554** | **4002.3108** | **0** |
| m6 | nbinom1(log) | ~Lineage + (1\|BlockParent.No) | ~Lineage | 11 | -1991.5783 | 4005.15652 | 2.84571927 |
| m8 | nbinom2(log) | ~Lineage + (1\|BlockParent.No) | ~1 | 8 | -2005.3249 | 4026.64989 | 24.3390901 |
| m9 | nbinom2(log) | ~Lineage + (1\|BlockParent.No) | ~Lineage | 11 | -2003.7478 | 4029.49561 | 27.1848088 |
| m11 | genpois(log) | ~Lineage + (1\|BlockParent.No) | ~1 | 8 | -2012.3771 | 4040.75414 | 38.4433407 |
| m12 | genpois(log) | ~Lineage + (1\|BlockParent.No) | ~Lineage | 11 | -2010.7999 | 4043.59986 | 41.2890601 |
| m4 | nbinom1(log) | ~Lineage + (1\|BlockParent.No) | ~0 | 7 | -2021.6215 | 4057.24309 | 54.9322962 |
| m10 | genpois(log) | ~Lineage + (1\|BlockParent.No) | ~0 | 7 | -2040.1965 | 4094.39302 | 92.0822222 |
| m7 | nbinom2(log) | ~Lineage + (1\|BlockParent.No) | ~0 | 7 | -2049.7511 | 4113.50215 | 111.191351 |
| m2 | poisson(log) | ~Lineage + (1\|BlockParent.No) | ~1 | 7 | -2219.4027 | 4452.80532 | 450.494525 |
| m3 | poisson(log) | ~Lineage + (1\|BlockParent.No) | ~Lineage | 10 | -2217.8255 | 4455.65104 | 453.340245 |

**Table S1I.** Full model selection for Age-Specific Reproduction in $F_2$.

**Significant zero-inflation was identified in a standard Poisson model (DHARMa zero-inflation test *p* = <0.001*).***

| Model | Family | Model formula | Zero-inflation formula | df | Loglik | AIC | Delta AIC |
|---|---|---|---|---|---|---|---|
| **m29** | **genpois(log)** | **~Lineage*Day2+ Lineage*Day + (1\|Block/Parent.No/Worm)** | **~Lineage+Day+Day2** | **38** | **-10977.29** | **22030.5802** | **0** |
| m12 | nbinom1(log) | ~Lineage*Day2+ Lineage*Day + (1\|Block/Parent.No/Worm) | ~Lineage+Day | 37 | -11004.152 | 22082.3032 | 51.7229103 |
| m11 | nbinom1(log) | ~Lineage*Day2+ Lineage*Day + (1\|Block/Parent.No/Worm) | ~Lineage | 36 | -11022.931 | 22117.8628 | 87.2825213 |
| m28 | genpois(log) | ~Lineage*Day2+ Lineage*Day + (1\|Block/Parent.No/Worm) | ~Lineage+Day | 37 | -11022.895 | 22119.7892 | 89.2089919 |
| m10 | nbinom1(log) | ~Lineage*Day2+ Lineage*Day + (1\|Block/Parent.No/Worm) | ~1 | 29 | -11032.309 | 22122.6182 | 92.0379679 |
| m27 | genpois(log) | ~Lineage*Day2+ Lineage*Day + (1\|Block/Parent.No/Worm) | ~Lineage | 36 | -11052.672 | 22177.3434 | 146.763127 |
| m26 | genpois(log) | ~Lineage*Day2+ Lineage*Day + (1\|Block/Parent.No/Worm) | ~1 | 29 | -11060.901 | 22179.8013 | 149.221055 |
| m22 | nbinom2(log) | ~Lineage*Day2+ Lineage*Day + (1\|Block/Parent.No/Worm) | ~Lineage*Day+Day2 | 45 | -11078.199 | 22246.3972 | 215.816989 |
| m24 | nbinom2(log) | ~Lineage*Day2+ Lineage*Day + (1\|Block/Parent.No/Worm) | ~Lineage*Day2+Lineage*Day | 52 | -11076.741 | 22257.4823 | 226.902026 |
| m9 | nbinom1(log) | ~Lineage*Day2+ Lineage*Day + (1\|Block/Parent.No/Worm) | ~0 | 28 | -11124.93 | 22305.8607 | 275.280499 |
| m21 | nbinom2(log) | ~Lineage*Day2+ Lineage*Day + (1\|Block/Parent.No/Worm) | ~Lineage+Day+Day2 | 38 | -11142.869 | 22361.7372 | 331.156952 |
| m20 | nbinom2(log) | ~Lineage*Day2+ Lineage*Day + (1\|Block/Parent.No/Worm) | ~Lineage+Day | 37 | -11170.077 | 22414.1534 | 383.573129 |
| m18 | nbinom2(log) | ~Lineage*Day2+ Lineage*Day + (1\|Block/Parent.No/Worm) | ~1 | 29 | -11413.659 | 22885.3171 | 854.736882 |
| m17 | nbinom2(log) | ~Lineage*Day2+ Lineage*Day + (1\|Block/Parent.No/Worm) | ~0 | 28 | -11473.411 | 23002.8215 | 972.241297 |
| m7 | poisson(log) | ~Lineage*Day2+ Lineage*Day + (1\|Block/Parent.No/Worm) | ~Lineage*Day2+Day | 44 | -16907.592 | 33903.1844 | 11872.6042 |
| m6 | poisson(log) | ~Lineage*Day2+ Lineage*Day + (1\|Block/Parent.No/Worm) | ~Lineage*Day+Day2 | 44 | -16922.385 | 33932.7702 | 11902.19 |
| m5 | poisson(log) | ~Lineage*Day2+ Lineage*Day + (1\|Block/Parent.No/Worm) | ~Lineage+Day+Day2 | 37 | -16965.526 | 34005.0513 | 11974.4711 |
| m4 | poisson(log) | ~Lineage*Day2+ Lineage*Day + (1\|Block/Parent.No/Worm) | ~Lineage+Day | 36 | -16970.88 | 34013.7594 | 11983.1791 |
| m3 | poisson(log) | ~Lineage*Day2+ Lineage*Day + (1\|Block/Parent.No/Worm) | ~Lineage | 35 | -17100.995 | 34271.9904 | 12241.4101 |
| m2 | poisson(log) | ~Lineage*Day2+ Lineage*Day + (1\|Block/Parent.No/Worm) | ~1 | 28 | -17116.351 | 34288.703 | 12258.1227 |
| m1 | poisson(log) | ~Lineage*Day2+ Lineage*Day + (1\|Block/Parent.No/Worm) | ~0 | 27 | -18412.597 | 36879.1947 | 14848.6144 |
| m8 | poisson(log) | ~Lineage*Day2+ Lineage*Day + (1\|Block/Parent.No/Worm) | ~Lineage*Day2+Lineage*Day | 51 | NA | NA | NA |
| m13 | nbinom1(log) | ~Lineage*Day2+ Lineage*Day + (1\|Block/Parent.No/Worm) | ~Lineage+Day+Day2 | 38 | NA | NA | NA |
| m14 | nbinom1(log) | ~Lineage*Day2+ Lineage*Day + (1\|Block/Parent.No/Worm) | ~Lineage*Day+Day2 | 45 | NA | NA | NA |
| m15 | nbinom1(log) | ~Lineage*Day2+ Lineage*Day + (1\|Block/Parent.No/Worm) | ~Lineage*Day2+Day | 45 | NA | NA | NA |
| m16 | nbinom1(log) | ~Lineage*Day2+ Lineage*Day + (1\|Block/Parent.No/Worm) | ~Lineage*Day2+Lineage*Day | 52 | NA | NA | NA |
| m19 | nbinom2(log) | ~Lineage*Day2+ Lineage*Day + (1\|Block/Parent.No/Worm) | ~Lineage | 36 | NA | NA | NA |
| m23 | nbinom2(log) | ~Lineage*Day2+ Lineage*Day + (1\|Block/Parent.No/Worm) | ~Lineage*Day2+Day | 45 | NA | NA | NA |
| m25 | genpois(log) | ~Lineage*Day2+ Lineage*Day + (1\|Block/Parent.No/Worm) | ~0 | 28 | NA | NA | NA |
| m30 | genpois(log) | ~Lineage*Day2+ Lineage*Day + (1\|Block/Parent.No/Worm) | ~Lineage*Day+Day2 | 45 | NA | NA | NA |
| m31 | genpois(log) | ~Lineage*Day2+ Lineage*Day + (1\|Block/Parent.No/Worm) | ~Lineage*Day2+Day | 45 | NA | NA | NA |
| m32 | genpois(log) | ~Lineage*Day2+ Lineage*Day + (1\|Block/Parent.No/Worm) | ~Lineage*Day2+Lineage*Day | 52 | NA | NA | NA |

**Table S1J.** Partial model selection for Age-Specific Reproduction in $F_2$ in temporary fasting conditions.

**Significant zero-inflation was identified in a standard Poisson model (DHARMa zero-inflation test *p* =<0.001*).***

| Model | Family | Model formula | Zero-inflation formula | df | Loglik | AIC | Delta AIC |
|---|---|---|---|---|---|---|---|
| m14 | nbinom1(log) | ~Lineage*Day2+ Lineage*Day + (1\|Block/Parent.No/Worm) | ~Lineage*Day+Day2 | 25 | -3327.4975 | 6704.99507 | 0 |
| **m28** | **genpois(log)** | **~Lineage*Day2+ Lineage*Day + (1\|Block/Parent.No/Worm)** | **~Lineage+Day** | **21** | **-3332.1203** | **6706.24053** | **1.245452** |

| | | | | | | | |
|---|---|---|---|---|---|---|---|
| m9 | nbinom1(log) | ~Lineage*Day2+ Lineage*Day + (1\|Block/Parent.No/Worm) | ~0 | 16 | -3366.3851 | 6764.77017 | 59.775091 |
| m25 | genpois(log) | ~Lineage*Day2+ Lineage*Day + (1\|Block/Parent.No/Worm) | ~0 | 16 | -3367.8791 | 6767.75828 | 62.763205 |
| m26 | genpois(log) | ~Lineage*Day2+ Lineage*Day + (1\|Block/Parent.No/Worm) | ~1 | 17 | -3367.1692 | 6768.33831 | 63.343237 |
| m11 | nbinom1(log) | ~Lineage*Day2+ Lineage*Day + (1\|Block/Parent.No/Worm) | ~Lineage | 20 | -3366.1052 | 6772.21035 | 67.215274 |
| m27 | genpois(log) | ~Lineage*Day2+ Lineage*Day + (1\|Block/Parent.No/Worm) | ~Lineage | 20 | -3366.2959 | 6772.59182 | 67.596749 |
| m12 | nbinom1(log) | ~Lineage*Day2+ Lineage*Day + (1\|Block/Parent.No/Worm) | ~Lineage+Day | 21 | -3366.3851 | 6774.77017 | 69.775091 |
| m18 | nbinom2(log) | ~Lineage*Day2+ Lineage*Day + (1\|Block/Parent.No/Worm) | ~1 | 17 | -3375.0965 | 6784.1931 | 79.198025 |
| m19 | nbinom2(log) | ~Lineage*Day2+ Lineage*Day + (1\|Block/Parent.No/Worm) | ~Lineage | 20 | -3374.0348 | 6788.0696 | 83.074529 |
| m17 | nbinom2(log) | ~Lineage*Day2+ Lineage*Day + (1\|Block/Parent.No/Worm) | ~0 | 16 | -3379.0077 | 6790.01536 | 85.020282 |
| m8 | poisson(log) | ~Lineage*Day2+ Lineage*Day + (1\|Block/Parent.No/Worm) | ~Lineage*Day2+Lineage*Day | 27 | -4845.2012 | 9744.40235 | 3039.40727 |
| m6 | poisson(log) | ~Lineage*Day2+ Lineage*Day + (1\|Block/Parent.No/Worm) | ~Lineage*Day+Day2 | 24 | -4853.5751 | 9755.15016 | 3050.15508 |
| m5 | poisson(log) | ~Lineage*Day2+ Lineage*Day + (1\|Block/Parent.No/Worm) | ~Lineage+Day+Day2 | 21 | -4857.3748 | 9756.74957 | 3051.7545 |
| m7 | poisson(log) | ~Lineage*Day2+ Lineage*Day + (1\|Block/Parent.No/Worm) | ~Lineage*Day2+Day | 24 | -4855.0843 | 9758.16863 | 3053.17355 |
| m4 | poisson(log) | ~Lineage*Day2+ Lineage*Day + (1\|Block/Parent.No/Worm) | ~Lineage+Day | 20 | -4879.5295 | 9799.05897 | 3094.06389 |
| m2 | poisson(log) | ~Lineage*Day2+ Lineage*Day + (1\|Block/Parent.No/Worm) | ~1 | 16 | -4987.1353 | 10006.2707 | 3301.27562 |
| m3 | poisson(log) | ~Lineage*Day2+ Lineage*Day + (1\|Block/Parent.No/Worm) | ~Lineage | 19 | -4984.6428 | 10007.2856 | 3302.2905 |
| m1 | poisson(log) | ~Lineage*Day2+ Lineage*Day + (1\|Block/Parent.No/Worm) | ~0 | 15 | -5376.8836 | 10783.7672 | 4078.77211 |
| m10 | nbinom1(log) | ~Lineage*Day2+ Lineage*Day + (1\|Block/Parent.No/Worm) | ~1 | 17 | NA | NA | NA |
| m13 | nbinom1(log) | ~Lineage*Day2+ Lineage*Day + (1\|Block/Parent.No/Worm) | ~Lineage+Day+Day2 | 22 | NA | NA | NA |
| m15 | nbinom1(log) | ~Lineage*Day2+ Lineage*Day + (1\|Block/Parent.No/Worm) | ~Lineage*Day2+Day | 25 | NA | NA | NA |
| m16 | nbinom1(log) | ~Lineage*Day2+ Lineage*Day + (1\|Block/Parent.No/Worm) | ~Lineage*Day2+Lineage*Day | 28 | NA | NA | NA |
| m20 | nbinom2(log) | ~Lineage*Day2+ Lineage*Day + (1\|Block/Parent.No/Worm) | ~Lineage+Day | 21 | NA | NA | NA |
| m21 | nbinom2(log) | ~Lineage*Day2+ Lineage*Day + (1\|Block/Parent.No/Worm) | ~Lineage+Day+Day2 | 22 | NA | NA | NA |
| m22 | nbinom2(log) | ~Lineage*Day2+ Lineage*Day + (1\|Block/Parent.No/Worm) | ~Lineage*Day+Day2 | 25 | NA | NA | NA |
| m23 | nbinom2(log) | ~Lineage*Day2+ Lineage*Day + (1\|Block/Parent.No/Worm) | ~Lineage*Day2+Day | 25 | NA | NA | NA |
| m24 | nbinom2(log) | ~Lineage*Day2+ Lineage*Day + (1\|Block/Parent.No/Worm) | ~Lineage*Day2+Lineage*Day | 28 | NA | NA | NA |
| m29 | genpois(log) | ~Lineage*Day2+ Lineage*Day + (1\|Block/Parent.No/Worm) | ~Lineage+Day+Day2 | 22 | NA | NA | NA |
| m30 | genpois(log) | ~Lineage*Day2+ Lineage*Day + (1\|Block/Parent.No/Worm) | ~Lineage*Day+Day2 | 25 | NA | NA | NA |
| m31 | genpois(log) | ~Lineage*Day2+ Lineage*Day + (1\|Block/Parent.No/Worm) | ~Lineage*Day2+Day | 25 | NA | NA | NA |
| m32 | genpois(log) | ~Lineage*Day2+ Lineage*Day + (1\|Block/Parent.No/Worm) | ~Lineage*Day2+Lineage*Day | 28 | NA | NA | NA |

**Table S1K.** Partial model selection for Age-Specific Reproduction in $F_2$ in *ad libitum* conditions.

**Significant zero-inflation was identified in a standard Poisson model (DHARMa zero-inflation test *p* = 0.024*).***

| Model | Family | Model formula | Zero-inflation formula | df | Loglik | AIC | Delta AIC |
|---|---|---|---|---|---|---|---|
| **m14** | **nbinom1(log)** | **~Lineage*Day2+ Lineage*Day + (1\|Block/Parent.No/Worm)** | **~Lineage*Day+Day2** | **25** | **-7583.0221** | **15216.0442** | **0** |
| m13 | nbinom1(log) | ~Lineage*Day2+ Lineage*Day + (1\|Block/Parent.No/Worm) | ~Lineage+Day+Day2 | 22 | -7587.0872 | 15218.1744 | 2.13019092 |
| m15 | nbinom1(log) | ~Lineage*Day2+ Lineage*Day + (1\|Block/Parent.No/Worm) | ~Lineage*Day2+Day | 25 | -7584.6992 | 15219.3984 | 3.35418149 |
| m30 | genpois(log) | ~Lineage*Day2+ Lineage*Day + (1\|Block/Parent.No/Worm) | ~Lineage*Day+Day2 | 25 | -7599.3587 | 15248.7174 | 32.6731963 |
| m29 | genpois(log) | ~Lineage*Day2+ Lineage*Day + (1\|Block/Parent.No/Worm) | ~Lineage+Day+Day2 | 22 | -7603.2158 | 15250.4316 | 34.3874679 |
| m12 | nbinom1(log) | ~Lineage*Day2+ Lineage*Day + (1\|Block/Parent.No/Worm) | ~Lineage+Day | 21 | -7633.8612 | 15309.7224 | 93.6782253 |

| | | | | | | | |
|---|---|---|---|---|---|---|---|
| m11 | nbinom1(log) | ~Lineage*Day2+ Lineage*Day + (1\|Block/Parent.No/Worm) | ~Lineage | 20 | -7645.1123 | 15330.2246 | 114.180421 |
| m10 | nbinom1(log) | ~Lineage*Day2+ Lineage*Day + (1\|Block/Parent.No/Worm) | ~1 | 17 | -7649.3689 | 15332.7378 | 116.693652 |
| m28 | genpois(log) | ~Lineage*Day2+ Lineage*Day + (1\|Block/Parent.No/Worm) | ~Lineage+Day | 21 | -7659.1767 | 15360.3533 | 144.309151 |
| m9 | nbinom1(log) | ~Lineage*Day2+ Lineage*Day + (1\|Block/Parent.No/Worm) | ~0 | 16 | -7727.6511 | 15487.3021 | 271.257951 |
| m22 | nbinom2(log) | ~Lineage*Day2+ Lineage*Day + (1\|Block/Parent.No/Worm) | ~Lineage*Day+Day2 | 25 | -7781.9675 | 15613.935 | 397.890793 |
| m24 | nbinom2(log) | ~Lineage*Day2+ Lineage*Day + (1\|Block/Parent.No/Worm) | ~Lineage*Day2+Lineage*Day | 28 | -7780.4434 | 15616.8867 | 400.842577 |
| m25 | genpois(log) | ~Lineage*Day2+ Lineage*Day + (1\|Block/Parent.No/Worm) | ~0 | 16 | -7795.4082 | 15622.8163 | 406.772155 |
| m20 | nbinom2(log) | ~Lineage*Day2+ Lineage*Day + (1\|Block/Parent.No/Worm) | ~Lineage+Day | 21 | -7854.2274 | 15750.4547 | 534.410564 |
| m19 | nbinom2(log) | ~Lineage*Day2+ Lineage*Day + (1\|Block/Parent.No/Worm) | ~Lineage | 20 | -8020.295 | 16080.5901 | 864.54592 |
| m18 | nbinom2(log) | ~Lineage*Day2+ Lineage*Day + (1\|Block/Parent.No/Worm) | ~1 | 17 | -8027.8207 | 16089.6413 | 873.597175 |
| m17 | nbinom2(log) | ~Lineage*Day2+ Lineage*Day + (1\|Block/Parent.No/Worm) | ~0 | 16 | -8083.5726 | 16199.1452 | 983.101027 |
| m6 | poisson(log) | ~Lineage*Day2+ Lineage*Day + (1\|Block/Parent.No/Worm) | ~Lineage*Day+Day2 | 24 | -11841.883 | 23731.7665 | 8515.72235 |
| m5 | poisson(log) | ~Lineage*Day2+ Lineage*Day + (1\|Block/Parent.No/Worm) | ~Lineage+Day+Day2 | 21 | -11845.029 | 23732.0583 | 8516.01411 |
| m7 | poisson(log) | ~Lineage*Day2+ Lineage*Day + (1\|Block/Parent.No/Worm) | ~Lineage*Day2+Day | 24 | -11843.239 | 23734.4772 | 8518.43299 |
| m4 | poisson(log) | ~Lineage*Day2+ Lineage*Day + (1\|Block/Parent.No/Worm) | ~Lineage+Day | 20 | -11935.203 | 23910.4056 | 8694.36144 |
| m3 | poisson(log) | ~Lineage*Day2+ Lineage*Day + (1\|Block/Parent.No/Worm) | ~Lineage | 19 | -11975.048 | 23988.095 | 8772.05087 |
| m2 | poisson(log) | ~Lineage*Day2+ Lineage*Day + (1\|Block/Parent.No/Worm) | ~1 | 16 | -11982.88 | 23997.7607 | 8781.71653 |
| m1 | poisson(log) | ~Lineage*Day2+ Lineage*Day + (1\|Block/Parent.No/Worm) | ~0 | 15 | -12943.984 | 25917.9677 | 10701.9235 |
| m8 | poisson(log) | ~Lineage*Day2+ Lineage*Day + (1\|Block/Parent.No/Worm) | ~Lineage*Day2+Lineage*Day | 27 | NA | NA | NA |
| m16 | nbinom1(log) | ~Lineage*Day2+ Lineage*Day + (1\|Block/Parent.No/Worm) | ~Lineage*Day2+Lineage*Day | 28 | NA | NA | NA |
| m21 | nbinom2(log) | ~Lineage*Day2+ Lineage*Day + (1\|Block/Parent.No/Worm) | ~Lineage+Day+Day2 | 22 | NA | NA | NA |
| m23 | nbinom2(log) | ~Lineage*Day2+ Lineage*Day + (1\|Block/Parent.No/Worm) | ~Lineage*Day2+Day | 25 | NA | NA | NA |
| m26 | genpois(log) | ~Lineage*Day2+ Lineage*Day + (1\|Block/Parent.No/Worm) | ~1 | 17 | NA | NA | NA |
| m27 | genpois(log) | ~Lineage*Day2+ Lineage*Day + (1\|Block/Parent.No/Worm) | ~Lineage | 20 | NA | NA | NA |
| m31 | genpois(log) | ~Lineage*Day2+ Lineage*Day + (1\|Block/Parent.No/Worm) | ~Lineage*Day2+Day | 25 | NA | NA | NA |
| m32 | genpois(log) | ~Lineage*Day2+ Lineage*Day + (1\|Block/Parent.No/Worm) | ~Lineage*Day2+Lineage*Day | 28 | NA | NA | NA |

**Table S1L.** Full model selection for Total Reproduction in $F_2$.

**Significant zero-inflation was identified in a standard Poisson model (DHARMa zero-inflation test *p* = <0.001*).***

| Model | Family | Model formula | Zero-inflation formula | df | Loglik | AIC | Delta AIC |
|---|---|---|---|---|---|---|---|
| **m5** | **nbinom1(log)** | **~Lineage + (1\|BlockParent.No)** | **~1** | **12** | **-2392.7782** | **4809.55641** | **0** |
| m11 | genpois(log) | ~Lineage + (1\|BlockParent.No) | ~1 | 12 | -2425.8698 | 4875.73967 | 66.1832616 |
| m12 | genpois(log) | ~Lineage + (1\|BlockParent.No) | ~Lineage | 19 | -2422.2244 | 4882.44883 | 72.8924158 |
| m4 | nbinom1(log) | ~Lineage + (1\|BlockParent.No) | ~0 | 11 | -2452.9112 | 4927.82239 | 118.265976 |
| m10 | genpois(log) | ~Lineage + (1\|BlockParent.No) | ~0 | 11 | -2497.3199 | 5016.63976 | 207.083346 |
| m8 | nbinom2(log) | ~Lineage + (1\|BlockParent.No) | ~1 | 12 | -2502.9092 | 5029.81833 | 220.261919 |
| m9 | nbinom2(log) | ~Lineage + (1\|BlockParent.No) | ~Lineage | 19 | -2499.2636 | 5036.52727 | 226.970863 |
| m7 | nbinom2(log) | ~Lineage + (1\|BlockParent.No) | ~0 | 11 | -2533.6315 | 5089.26297 | 279.706563 |
| m2 | poisson(log) | ~Lineage + (1\|BlockParent.No) | ~1 | 11 | -3933.566 | 7889.13204 | 3079.57563 |
| m3 | poisson(log) | ~Lineage + (1\|BlockParent.No) | ~Lineage | 18 | -3929.9206 | 7895.84119 | 3086.28478 |

| | | | | | | | |
|---|---|---|---|---|---|---|---|
| m6 | nbinom1(log) | ~Lineage + (1\|BlockParent.No) | ~Lineage | 19 | NA | NA | NA |

**Table S1M.** Partial model selection for Total Reproduction in $F_2$ in temporary fasting conditions.

**No significant zero-inflation was identified in a standard Poisson model (DHARMa zero-inflation test *p* = 1*).***

| Model | Family | Model formula | df | Loglik | AIC | Delta AIC |
|---|---|---|---|---|---|---|
| **m4** | **nbinom1(log)** | **~Lineage + (1\|BlockParent.No)** | **7** | **-762.38712** | **1538.77424** | **0** |
| m7 | nbinom2(log) | ~Lineage + (1\|BlockParent.No) | 7 | -767.24987 | 1548.49974 | 9.72549365 |
| m10 | genpois(log) | ~Lineage + (1\|BlockParent.No) | 7 | -768.35415 | 1550.70831 | 11.9340639 |
| m0 | poisson(log) | ~Lineage + (1\|BlockParent.No) | 6 | -1031.0288 | 2074.05759 | 535.283347 |

**Table S1N.** Partial model selection for Total Reproduction in $F_2$ in *ad libitum* conditions.

**Significant zero-inflation was identified in a standard Poisson model (DHARMa zero-inflation test *p* = <0.001*).***

| Model | Family | Model formula | Zero-inflation formula | df | Loglik | AIC | Delta AIC |
|---|---|---|---|---|---|---|---|
| **m5** | **nbinom1(log)** | **~Lineage + (1\|BlockParent.No)** | **~1** | **8** | **-1505.974** | **3027.948** | **0** |
| m6 | nbinom1(log) | ~Lineage + (1\|BlockParent.No) | ~Lineage | 11 | -1503.0955 | 3028.191 | 0.2430013 |
| m11 | genpois(log) | ~Lineage + (1\|BlockParent.No) | ~1 | 8 | -1508.128 | 3032.25605 | 4.30804751 |
| m12 | genpois(log) | ~Lineage + (1\|BlockParent.No) | ~Lineage | 11 | -1505.2495 | 3032.49905 | 4.55104859 |
| m8 | nbinom2(log) | ~Lineage + (1\|BlockParent.No) | ~1 | 8 | -1509.8771 | 3035.75417 | 7.80617217 |
| m9 | nbinom2(log) | ~Lineage + (1\|BlockParent.No) | ~Lineage | 11 | -1506.9986 | 3035.99717 | 8.04917297 |
| m2 | poisson(log) | ~Lineage + (1\|BlockParent.No) | ~1 | 7 | -1534.2992 | 3082.59837 | 54.6503721 |
| m3 | poisson(log) | ~Lineage + (1\|BlockParent.No) | ~Lineage | 10 | -1531.4207 | 3082.84137 | 54.8933732 |
| m4 | nbinom1(log) | ~Lineage + (1\|BlockParent.No) | ~0 | 7 | -1635.4709 | 3284.94185 | 256.993848 |
| m7 | nbinom2(log) | ~Lineage + (1\|BlockParent.No) | ~0 | 7 | -1647.3375 | 3308.67494 | 280.726943 |
| m10 | genpois(log) | ~Lineage + (1\|BlockParent.No) | ~0 | 7 | -1649.3793 | 3312.75862 | 284.810621 |

**Table S1O.** Full model selection for Age-Specific Reproduction in $F_3$.

**No significant zero-inflation was identified in a standard Poisson model (DHARMa zero-inflation test *p* = 0.256*).***

| Model | Family | Model formula | df | Loglik | AIC | Delta AIC |
|---|---|---|---|---|---|---|
| **m9** | **nbinom1(log)** | **~Lineage*Day2+ Lineage*Day + (1\|Block/Parent.No/Worm)** | **28** | **-10498.638** | **21053.2751** | **0** |
| m25 | genpois(log) | ~Lineage*Day2+ Lineage*Day + (1\|Block/Parent.No/Worm) | 28 | -10551.724 | 21159.4481 | 106.172987 |
| m17 | nbinom2(log) | ~Lineage*Day2+ Lineage*Day + (1\|Block/Parent.No/Worm) | 28 | -11129.903 | 22315.8061 | 1262.53101 |
| m1 | poisson(log) | ~Lineage*Day2+ Lineage*Day + (1\|Block/Parent.No/Worm) | 27 | -16813.737 | 33681.4745 | 12628.1994 |

**Table S1P.** Partial model selection for Age-Specific Reproduction in $F_3$ in temporary fasting conditions.

**Significant zero-inflation was identified in a standard Poisson model (DHARMa zero-inflation test *p* =<0.001*).***

| Model | Family | Model formula | Zero-inflation formula | df | Loglik | AIC | Delta AIC |
|---|---|---|---|---|---|---|---|
| **m12** | **nbinom1(log)** | **~Lineage*Day2+ Lineage*Day + (1\|Block/Parent.No/Worm)** | **~Lineage+Day** | **21** | **-2648.4417** | **5338.88336** | **0** |
| m28 | genpois(log) | ~Lineage*Day2+ Lineage*Day + (1\|Block/Parent.No/Worm) | ~Lineage+Day | 21 | -2656.6108 | 5355.22162 | 16.3382683 |
| m10 | nbinom1(log) | ~Lineage*Day2+ Lineage*Day + (1\|Block/Parent.No/Worm) | ~1 | 17 | -2671.3951 | 5376.79011 | 37.9067536 |
| m9 | nbinom1(log) | ~Lineage*Day2+ Lineage*Day + (1\|Block/Parent.No/Worm) | ~0 | 16 | -2672.9952 | 5377.99048 | 39.1071242 |
| m11 | nbinom1(log) | ~Lineage*Day2+ Lineage*Day + (1\|Block/Parent.No/Worm) | ~Lineage | 20 | -2669.6025 | 5379.2051 | 40.3217433 |
| m13 | nbinom1(log) | ~Lineage*Day2+ Lineage*Day + (1\|Block/Parent.No/Worm) | ~Lineage+Day+Day2 | 22 | -2670.0839 | 5384.16778 | 45.2844218 |
| m26 | genpois(log) | ~Lineage*Day2+ Lineage*Day + (1\|Block/Parent.No/Worm) | ~1 | 17 | -2683.2469 | 5400.49373 | 61.6103778 |
| m27 | genpois(log) | ~Lineage*Day2+ Lineage*Day + (1\|Block/Parent.No/Worm) | ~Lineage | 20 | -2680.6053 | 5401.21059 | 62.3272345 |
| m25 | genpois(log) | ~Lineage*Day2+ Lineage*Day + (1\|Block/Parent.No/Worm) | ~0 | 16 | -2686.2329 | 5404.46575 | 65.5823964 |
| m18 | nbinom2(log) | ~Lineage*Day2+ Lineage*Day + (1\|Block/Parent.No/Worm) | ~1 | 17 | -2779.4151 | 5592.83016 | 253.946799 |
| m19 | nbinom2(log) | ~Lineage*Day2+ Lineage*Day + (1\|Block/Parent.No/Worm) | ~Lineage | 20 | -2777.0877 | 5594.17538 | 255.292027 |
| m17 | nbinom2(log) | ~Lineage*Day2+ Lineage*Day + (1\|Block/Parent.No/Worm) | ~0 | 16 | -2788.6681 | 5609.33622 | 270.452859 |
| m6 | poisson(log) | ~Lineage*Day2+ Lineage*Day + (1\|Block/Parent.No/Worm) | ~Lineage*Day+Day2 | 24 | -3689.0542 | 7426.10842 | 2087.22507 |
| m8 | poisson(log) | ~Lineage*Day2+ Lineage*Day + (1\|Block/Parent.No/Worm) | ~Lineage*Day2+Lineage*Day | 27 | -3688.0553 | 7430.11054 | 2091.22718 |
| m7 | poisson(log) | ~Lineage*Day2+ Lineage*Day + (1\|Block/Parent.No/Worm) | ~Lineage*Day2+Day | 24 | -3695.024 | 7438.04794 | 2099.16459 |
| m5 | poisson(log) | ~Lineage*Day2+ Lineage*Day + (1\|Block/Parent.No/Worm) | ~Lineage+Day+Day2 | 21 | -3715.0561 | 7472.11222 | 2133.22887 |
| m3 | poisson(log) | ~Lineage*Day2+ Lineage*Day + (1\|Block/Parent.No/Worm) | ~Lineage | 19 | -3767.149 | 7572.29809 | 2233.41473 |
| m2 | poisson(log) | ~Lineage*Day2+ Lineage*Day + (1\|Block/Parent.No/Worm) | ~1 | 16 | -3771.4215 | 7574.84296 | 2235.9596 |
| m1 | poisson(log) | ~Lineage*Day2+ Lineage*Day + (1\|Block/Parent.No/Worm) | ~0 | 15 | -4028.3125 | 8086.62495 | 2747.74159 |
| m4 | poisson(log) | ~Lineage*Day2+ Lineage*Day + (1\|Block/Parent.No/Worm) | ~Lineage+Day | 20 | NA | NA | NA |
| m14 | nbinom1(log) | ~Lineage*Day2+ Lineage*Day + (1\|Block/Parent.No/Worm) | ~Lineage*Day+Day2 | 25 | NA | NA | NA |
| m15 | nbinom1(log) | ~Lineage*Day2+ Lineage*Day + (1\|Block/Parent.No/Worm) | ~Lineage*Day2+Day | 25 | NA | NA | NA |
| m16 | nbinom1(log) | ~Lineage*Day2+ Lineage*Day + (1\|Block/Parent.No/Worm) | ~Lineage*Day2+Lineage*Day | 28 | NA | NA | NA |
| m20 | nbinom2(log) | ~Lineage*Day2+ Lineage*Day + (1\|Block/Parent.No/Worm) | ~Lineage+Day | 21 | NA | NA | NA |
| m21 | nbinom2(log) | ~Lineage*Day2+ Lineage*Day + (1\|Block/Parent.No/Worm) | ~Lineage+Day+Day2 | 22 | NA | NA | NA |
| m22 | nbinom2(log) | ~Lineage*Day2+ Lineage*Day + (1\|Block/Parent.No/Worm) | ~Lineage*Day+Day2 | 25 | NA | NA | NA |
| m23 | nbinom2(log) | ~Lineage*Day2+ Lineage*Day + (1\|Block/Parent.No/Worm) | ~Lineage*Day2+Day | 25 | NA | NA | NA |
| m24 | nbinom2(log) | ~Lineage*Day2+ Lineage*Day + (1\|Block/Parent.No/Worm) | ~Lineage*Day2+Lineage*Day | 28 | NA | NA | NA |
| m29 | genpois(log) | ~Lineage*Day2+ Lineage*Day + (1\|Block/Parent.No/Worm) | ~Lineage+Day+Day2 | 22 | NA | NA | NA |
| m30 | genpois(log) | ~Lineage*Day2+ Lineage*Day + (1\|Block/Parent.No/Worm) | ~Lineage*Day+Day2 | 25 | NA | NA | NA |
| m31 | genpois(log) | ~Lineage*Day2+ Lineage*Day + (1\|Block/Parent.No/Worm) | ~Lineage*Day2+Day | 25 | NA | NA | NA |
| m32 | genpois(log) | ~Lineage*Day2+ Lineage*Day + (1\|Block/Parent.No/Worm) | ~Lineage*Day2+Lineage*Day | 28 | NA | NA | NA |

**Table S1Q.** Partial model selection for Age-Specific Reproduction in $F_3$ in *ad libitum* conditions.

**Significant zero-inflation was identified in a standard Poisson model (DHARMa zero-inflation test *p* =<0.001*).***

| Model | Family | Model formula | Zero-inflation formula | df | Loglik | AIC | Delta AIC |
|---|---|---|---|---|---|---|---|
| **m10** | **nbinom1(log)** | **~Lineage*Day2+ Lineage*Day + (1\|Block/Parent.No/Worm)** | **~1** | **17** | **-7703.9204** | **15441.8409** | **0** |
| m13 | nbinom1(log) | ~Lineage*Day2+ Lineage*Day + (1\|Block/Parent.No/Worm) | ~Lineage+Day+Day2 | 22 | -7699.7625 | 15443.5251 | 1.68416721 |
| m12 | nbinom1(log) | ~Lineage*Day2+ Lineage*Day + (1\|Block/Parent.No/Worm) | ~Lineage+Day | 21 | -7701.6749 | 15445.3498 | 3.50892234 |
| m14 | nbinom1(log) | ~Lineage*Day2+ Lineage*Day + (1\|Block/Parent.No/Worm) | ~Lineage*Day+Day2 | 25 | -7698.0578 | 15446.1156 | 4.27466969 |
| m16 | nbinom1(log) | ~Lineage*Day2+ Lineage*Day + (1\|Block/Parent.No/Worm) | ~Lineage*Day2+Lineage*Day | 28 | -7697.0666 | 15450.1332 | 8.29234444 |
| m15 | nbinom1(log) | ~Lineage*Day2+ Lineage*Day + (1\|Block/Parent.No/Worm) | ~Lineage*Day2+Day | 25 | -7707.0801 | 15464.1602 | 22.3193261 |
| m29 | genpois(log) | ~Lineage*Day2+ Lineage*Day + (1\|Block/Parent.No/Worm) | ~Lineage+Day+Day2 | 22 | -7728.0544 | 15500.1089 | 58.2679912 |
| m26 | genpois(log) | ~Lineage*Day2+ Lineage*Day + (1\|Block/Parent.No/Worm) | ~1 | 17 | -7733.5914 | 15501.1828 | 59.341864 |
| m30 | genpois(log) | ~Lineage*Day2+ Lineage*Day + (1\|Block/Parent.No/Worm) | ~Lineage*Day+Day2 | 25 | -7726.3182 | 15502.6365 | 60.7955977 |
| m27 | genpois(log) | ~Lineage*Day2+ Lineage*Day + (1\|Block/Parent.No/Worm) | ~Lineage | 20 | -7731.7365 | 15503.4729 | 61.6320047 |
| m28 | genpois(log) | ~Lineage*Day2+ Lineage*Day + (1\|Block/Parent.No/Worm) | ~Lineage+Day | 21 | -7730.9848 | 15503.9697 | 62.1287904 |
| m9 | nbinom1(log) | ~Lineage*Day2+ Lineage*Day + (1\|Block/Parent.No/Worm) | ~0 | 16 | -7806.6423 | 15645.2846 | 203.443677 |
| m25 | genpois(log) | ~Lineage*Day2+ Lineage*Day + (1\|Block/Parent.No/Worm) | ~0 | 16 | -7850.4518 | 15732.9035 | 291.062637 |
| m24 | nbinom2(log) | ~Lineage*Day2+ Lineage*Day + (1\|Block/Parent.No/Worm) | ~Lineage*Day2+Lineage*Day | 28 | -8130.9515 | 16317.903 | 876.0621 |
| m20 | nbinom2(log) | ~Lineage*Day2+ Lineage*Day + (1\|Block/Parent.No/Worm) | ~Lineage+Day | 21 | -8154.0113 | 16350.0225 | 908.181621 |
| m18 | nbinom2(log) | ~Lineage*Day2+ Lineage*Day + (1\|Block/Parent.No/Worm) | ~1 | 17 | -8308.3911 | 16650.7822 | 1208.94135 |
| m19 | nbinom2(log) | ~Lineage*Day2+ Lineage*Day + (1\|Block/Parent.No/Worm) | ~Lineage | 20 | -8306.1279 | 16652.2558 | 1210.41487 |
| m17 | nbinom2(log) | ~Lineage*Day2+ Lineage*Day + (1\|Block/Parent.No/Worm) | ~0 | 16 | -8331.9483 | 16695.8967 | 1254.05577 |
| m5 | poisson(log) | ~Lineage*Day2+ Lineage*Day + (1\|Block/Parent.No/Worm) | ~Lineage+Day+Day2 | 21 | -12155 | 24351.9998 | 8910.1589 |
| m6 | poisson(log) | ~Lineage*Day2+ Lineage*Day + (1\|Block/Parent.No/Worm) | ~Lineage*Day+Day2 | 24 | -12154.584 | 24357.1671 | 8915.32622 |
| m7 | poisson(log) | ~Lineage*Day2+ Lineage*Day + (1\|Block/Parent.No/Worm) | ~Lineage*Day2+Day | 24 | -12154.744 | 24357.4874 | 8915.64655 |
| m8 | poisson(log) | ~Lineage*Day2+ Lineage*Day + (1\|Block/Parent.No/Worm) | ~Lineage*Day2+Lineage*Day | 27 | -12152.697 | 24359.3933 | 8917.55243 |
| m4 | poisson(log) | ~Lineage*Day2+ Lineage*Day + (1\|Block/Parent.No/Worm) | ~Lineage+Day | 20 | -12176.676 | 24393.3518 | 8951.51089 |
| m2 | poisson(log) | ~Lineage*Day2+ Lineage*Day + (1\|Block/Parent.No/Worm) | ~1 | 16 | -12197.262 | 24426.5232 | 8984.6823 |
| m3 | poisson(log) | ~Lineage*Day2+ Lineage*Day + (1\|Block/Parent.No/Worm) | ~Lineage | 19 | -12195.746 | 24429.4925 | 8987.65156 |
| m1 | poisson(log) | ~Lineage*Day2+ Lineage*Day + (1\|Block/Parent.No/Worm) | ~0 | 15 | -12767.324 | 25564.6481 | 10122.8072 |
| m11 | nbinom1(log) | ~Lineage*Day2+ Lineage*Day + (1\|Block/Parent.No/Worm) | ~Lineage | 20 | NA | NA | NA |
| m21 | nbinom2(log) | ~Lineage*Day2+ Lineage*Day + (1\|Block/Parent.No/Worm) | ~Lineage+Day+Day2 | 22 | NA | NA | NA |
| m22 | nbinom2(log) | ~Lineage*Day2+ Lineage*Day + (1\|Block/Parent.No/Worm) | ~Lineage*Day+Day2 | 25 | NA | NA | NA |
| m23 | nbinom2(log) | ~Lineage*Day2+ Lineage*Day + (1\|Block/Parent.No/Worm) | ~Lineage*Day2+Day | 25 | NA | NA | NA |
| m31 | genpois(log) | ~Lineage*Day2+ Lineage*Day + (1\|Block/Parent.No/Worm) | ~Lineage*Day2+Day | 25 | NA | NA | NA |
| m32 | genpois(log) | ~Lineage*Day2+ Lineage*Day + (1\|Block/Parent.No/Worm) | ~Lineage*Day2+Lineage*Day | 28 | NA | NA | NA |

**Table S1R.** Full model selection for Total Reproduction in $F_3$.

**Significant zero-inflation was identified in a standard Poisson model (DHARMa zero-inflation test *p* = <0.001*).***

| Model | Family | Model formula | Zero-inflation formula | df | Loglik | AIC | Delta AIC |
|---|---|---|---|---|---|---|---|

| | | | | | | | |
|---|---|---|---|---|---|---|---|
| **m6** | **nbinom1(log)** | **~Lineage + (1\|BlockParent.No)** | **~Lineage** | **19** | **-2293.6246** | **4625.24916** | **0** |
| m5 | nbinom1(log) | ~Lineage + (1\|BlockParent.No) | ~1 | 12 | -2303.8532 | 4631.70632 | 6.45715969 |
| m12 | genpois(log) | ~Lineage + (1\|BlockParent.No) | ~Lineage | 19 | -2336.8365 | 4711.67308 | 86.4239196 |
| m11 | genpois(log) | ~Lineage + (1\|BlockParent.No) | ~1 | 12 | -2347.0651 | 4718.13024 | 92.8810809 |
| m9 | nbinom2(log) | ~Lineage + (1\|BlockParent.No) | ~Lineage | 19 | -2359.6722 | 4757.34439 | 132.095232 |
| m8 | nbinom2(log) | ~Lineage + (1\|BlockParent.No) | ~1 | 12 | -2369.9008 | 4763.80159 | 138.552436 |
| m4 | nbinom1(log) | ~Lineage + (1\|BlockParent.No) | ~0 | 11 | -2379.4309 | 4780.86184 | 155.612685 |
| m10 | genpois(log) | ~Lineage + (1\|BlockParent.No) | ~0 | 11 | -2445.8454 | 4913.69071 | 288.441549 |
| m7 | nbinom2(log) | ~Lineage + (1\|BlockParent.No) | ~0 | 11 | -2461.4975 | 4944.99501 | 319.745853 |
| m3 | poisson(log) | ~Lineage + (1\|BlockParent.No) | ~Lineage | 18 | -3422.0048 | 6880.00963 | 2254.76047 |
| m2 | poisson(log) | ~Lineage + (1\|BlockParent.No) | ~1 | 11 | -3432.2334 | 6886.46679 | 2261.21763 |

**Table S1S.** Partial model selection for Total Reproduction in $F_3$ in temporary fasting conditions.

**Significant zero-inflation was identified in a standard Poisson model (DHARMa zero-inflation test *p* = <0.001*).***

| Model | Family | Model formula | Zero-inflation formula | df | Loglik | AIC | Delta AIC |
|---|---|---|---|---|---|---|---|
| **m6** | **nbinom1(log)** | **~Lineage + (1\|BlockParent.No)** | **~Lineage** | **11** | **-623.08241** | **1268.16481** | **0** |
| m5 | nbinom1(log) | ~Lineage + (1\|BlockParent.No) | ~1 | 8 | -627.71156 | 1271.42312 | 3.25830392 |
| m12 | genpois(log) | ~Lineage + (1\|BlockParent.No) | ~Lineage | 11 | -630.9575 | 1283.91499 | 15.7501795 |
| m11 | genpois(log) | ~Lineage + (1\|BlockParent.No) | ~1 | 8 | -635.58668 | 1287.17337 | 19.0085535 |
| m4 | nbinom1(log) | ~Lineage + (1\|BlockParent.No) | ~0 | 7 | -646.44612 | 1306.89224 | 38.7274247 |
| m7 | nbinom2(log) | ~Lineage + (1\|BlockParent.No) | ~0 | 7 | -654.65513 | 1323.31026 | 55.1454519 |
| m10 | genpois(log) | ~Lineage + (1\|BlockParent.No) | ~0 | 7 | -661.7532 | 1337.50641 | 69.3415936 |
| m3 | poisson(log) | ~Lineage + (1\|BlockParent.No) | ~Lineage | 10 | -762.41856 | 1544.83711 | 276.6723 |
| m2 | poisson(log) | ~Lineage + (1\|BlockParent.No) | ~1 | 7 | -767.04774 | 1548.09549 | 279.930674 |
| m8 | nbinom2(log) | ~Lineage + (1\|BlockParent.No) | ~1 | 8 | NA | NA | NA |
| m9 | nbinom2(log) | ~Lineage + (1\|BlockParent.No) | ~Lineage | 11 | NA | NA | NA |

**Table S1T.** Partial model selection for Total Reproduction in $F_3$ in *ad libitum* conditions.

**Significant zero-inflation was identified in a standard Poisson model (DHARMa zero-inflation test *p* = <0.001*).***

| Model | Family | Model formula | Zero-inflation formula | df | Loglik | AIC | Delta AIC |
|---|---|---|---|---|---|---|---|
| **m5** | **nbinom1(log)** | **~Lineage + (1\|BlockParent.No)** | **~1** | **8** | **-1588.1106** | **3192.22114** | **0** |
| m6 | nbinom1(log) | ~Lineage + (1\|BlockParent.No) | ~Lineage | 11 | -1586.6632 | 3195.32644 | 3.10529994 |
| m8 | nbinom2(log) | ~Lineage + (1\|BlockParent.No) | ~1 | 8 | -1597.6889 | 3211.37783 | 19.1566907 |
| m9 | nbinom2(log) | ~Lineage + (1\|BlockParent.No) | ~Lineage | 11 | -1596.2416 | 3214.48313 | 22.2619903 |
| m11 | genpois(log) | ~Lineage + (1\|BlockParent.No) | ~1 | 8 | -1600.4227 | 3216.84538 | 24.6242415 |
| m12 | genpois(log) | ~Lineage + (1\|BlockParent.No) | ~Lineage | 11 | -1598.9753 | 3219.95068 | 27.7295415 |

| | | | | | | | |
|---|---|---|---|---|---|---|---|
| m4 | nbinom1(log) | ~Lineage + (1\|BlockParent.No) | ~0 | 7 | -1643.3362 | 3300.67238 | 108.451238 |
| m7 | nbinom2(log) | ~Lineage + (1\|BlockParent.No) | ~0 | 7 | -1657.8165 | 3329.633 | 137.411862 |
| m10 | genpois(log) | ~Lineage + (1\|BlockParent.No) | ~0 | 7 | -1664.9061 | 3343.81214 | 151.590999 |
| m2 | poisson(log) | ~Lineage + (1\|BlockParent.No) | ~1 | 7 | -1705.149 | 3424.29807 | 232.07693 |
| m3 | poisson(log) | ~Lineage + (1\|BlockParent.No) | ~Lineage | 10 | -1703.7017 | 3427.40337 | 235.18223 |

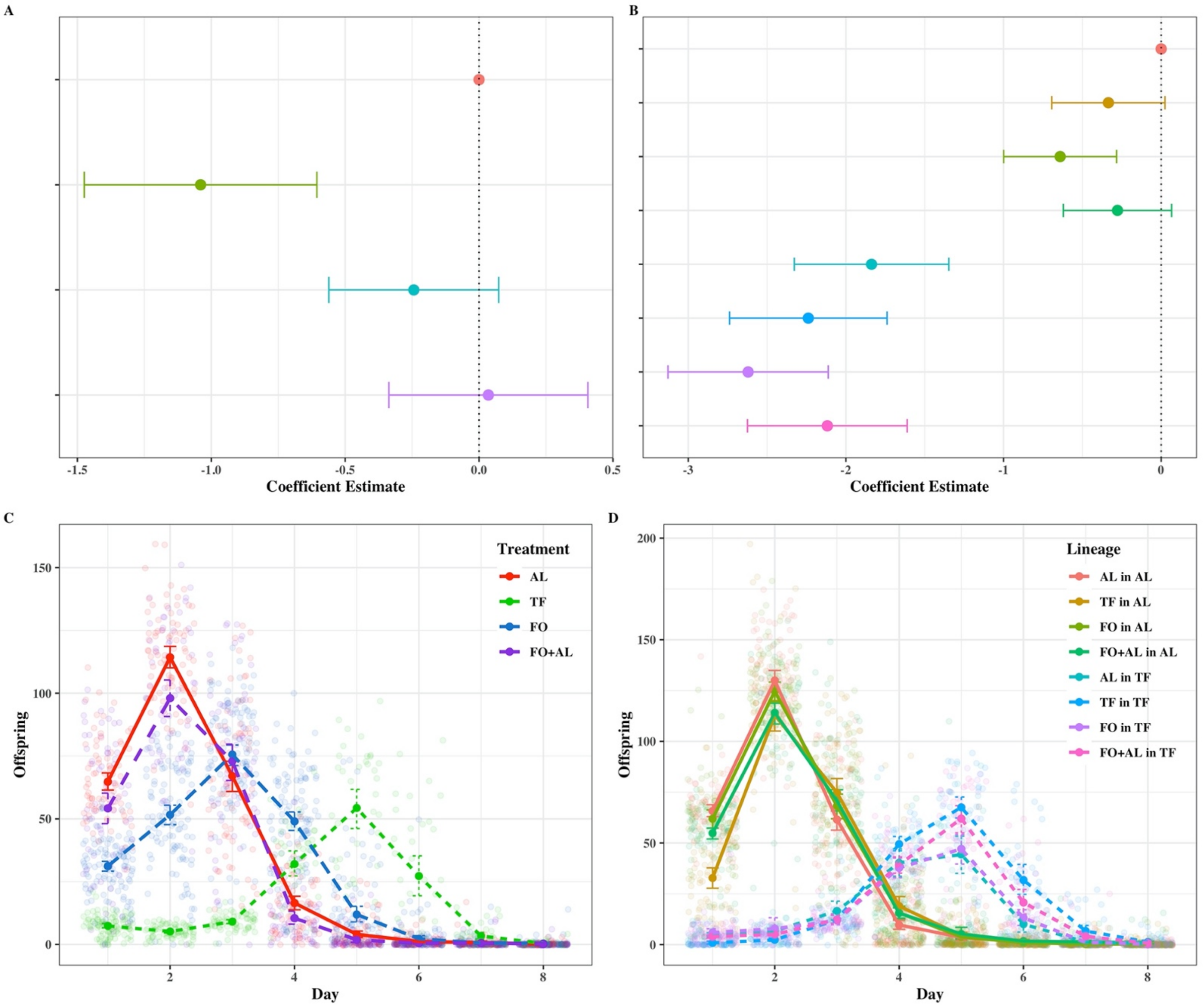

**Fig. S1A-D** Plots of survival (A-B) and reproduction (C-D) for $P_0$ (A, C) and $F_1$ (B, D) **without Block 1**. Colours represent differing dietary treatments for each generation. For Graph D the lineages are given as $P_0$ in F1. A-B) Points represent coefficients from a mixed effects cox model with 95% confidence intervals; C-D) Points represent mean values with 95% confidence intervals.

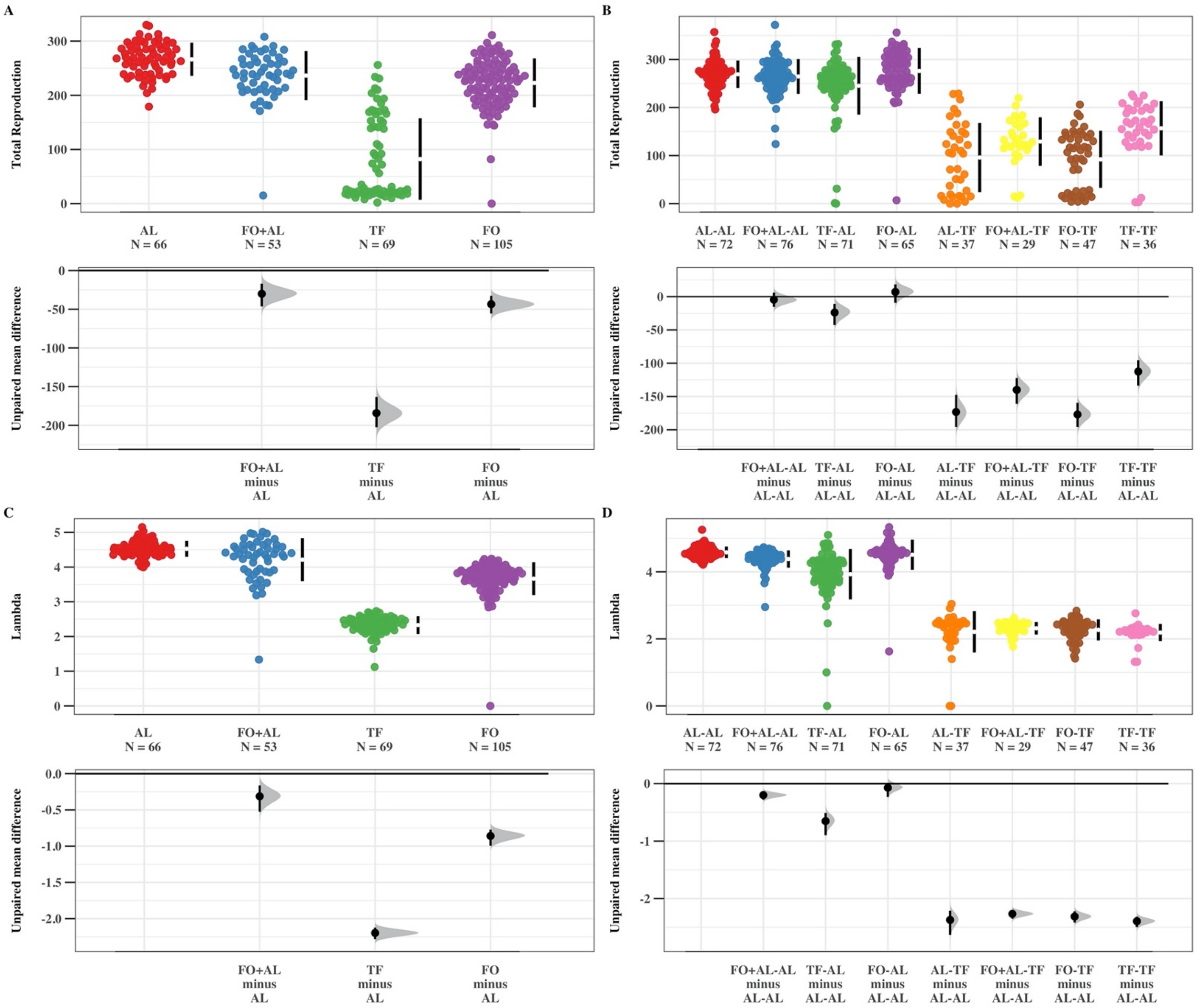

**Fig. S2A-D** Plots of LRS (A-B) and lambda (C-D) for $P_0$ (A, C) and $F_1$ (B, D) **without Block 1**. Colours represent differing dietary treatments for each generation. For Graphs B and D, the lineages are given as $P_0$ -$F_1$. Bootstrapped mean comparison between treatments, showing mean values with 95% confidence intervals in both top and bottom panels

**Table S2A.** Full model output from the Mixed Effects Cox Proportional Hazard Model for survival in $P_0$. In all cases, the treatment not shown in the output is the reference factor. Treatments denoted in bold are significantly different from zero.

| Covariate | Estimate | SE | *z* value | *p* value | Low 95% CI | High 95% CI |
|---|---|---|---|---|---|---|
| **TF** | **-1.104** | **0.197** | **-5.591** | **<0.001** | **-1.490** | **-0.717** |
| **FO** | **-0.356** | **0.141** | **-2.519** | **0.018** | **-0.633** | **-0.0791** |
| FO+AL | 0.295 | 0.166 | 1.770 | 0.077 | -0.0316 | 0.621 |

**Table S2B.** Full model output from the best identified model for Age-Specific Reproduction in $P_0$. Covariates denoted in bold are significant to $\alpha = 0.05$.

| Model | Covariate | Estimate | SE | *z* value | *p* value |
|---|---|---|---|---|---|
| Conditional | **(Intercept)** | **3.669** | **0.082** | **44.610** | **<0.001** |
| | **TF** | **-3.080** | **0.166** | **-18.520** | **<0.001** |
| | **FO** | **-1.455** | **0.113** | **-12.850** | **<0.001** |
| | **FO+AL** | **-0.789** | **0.125** | **-6.310** | **<0.001** |
| | **Day2** | **-0.255** | **0.012** | **-20.750** | **<0.001** |
| | **Day** | **0.934** | **0.065** | **14.310** | **<0.001** |
| | **TF:Day2** | **0.132** | **0.016** | **8.280** | **<0.001** |
| | FO:Day2 | -0.024 | 0.015 | -1.620 | 0.1046 |
| | **FO+AL:Day2** | **-0.118** | **0.020** | **-5.750** | **<0.001** |
| | TF:Day | 0.185 | 0.106 | 1.750 | 0.0804 |
| | **FO:Day** | **0.563** | **0.086** | **6.520** | **<0.001** |
| | **FO+AL:Day** | **0.616** | **0.108** | **5.700** | **<0.001** |

| | | | | | |
|---|---|---|---|---|---|
| Zero-Inflation | (Intercept) | -178.991 | 13361.764 | -0.013 | 0.989 |
| | TF | 155.269 | 13361.674 | 0.012 | 0.991 |
| | FO | 170.561 | 13361.770 | 0.013 | 0.990 |
| | FO+AL | 171.094 | 13361.753 | 0.013 | 0.990 |
| | Day2 | -8.373 | 668.037 | -0.013 | 0.990 |
| | Day | 77.358 | 6012.628 | 0.013 | 0.990 |
| | TF:Day2 | 8.054 | 668.034 | 0.012 | 0.990 |
| | FO:Day2 | 8.223 | 668.037 | 0.012 | 0.990 |
| | FO+AL:Day2 | 8.136 | 668.036 | 0.012 | 0.990 |
| | TF:Day | -71.744 | 6012.596 | -0.012 | 0.990 |
| | FO:Day | -75.467 | 6012.632 | -0.013 | 0.990 |
| | FO+AL:Day | -75.087 | 6012.624 | -0.012 | 0.990 |

**Table S2C.** Full model output from the best identified model for Total Reproduction in $P_0$.

| Model | Covariate | Estimate | SE | *z* value | *p* value |
|---|---|---|---|---|---|
| Conditional | **(Intercept)** | **5.602** | **0.058** | **96.350** | **<0.001** |
| | **TF** | **-2.192** | **0.074** | **-29.670** | **<0.001** |
| | **FO** | **-0.206** | **0.039** | **-5.260** | **<0.001** |
| | **FO+AL** | **-0.148** | **0.044** | **-3.400** | **<0.001** |
| Zero-Inflation | **(Intercept)** | **-5.223** | **0.759** | **-6.882** | **<0.001** |

**Table S2D.** Full model output from the mixed effect model for individual fitness ($\lambda$) in $P_0$.

| Covariate | Estimate | SE | *z* value | *p* value |
|---|---|---|---|---|
| **(Intercept)** | **4.486** | **0.088** | **51.190** | **<0.001** |
| **TF** | **-2.419** | **0.057** | **-42.220** | **<0.001** |
| **FO** | **-0.879** | **0.057** | **-15.470** | **<0.001** |
| **FO+AL** | **-0.267** | **0.062** | **-4.280** | **<0.001** |

**Table S2E.** Full model output from the Mixed Effects Cox Proportional Hazard Model for survival in $F_1$.

| $P_0$ | $F_1$ | Estimate | SE | *z* value | *p* value | Low 95% CI | High 95% CI |
|---|---|---|---|---|---|---|---|
| TF | AL | -0.330 | 0.188 | -1.753 | 0.080 | -0.698 | 0.039 |
| **FO** | **AL** | **-0.691** | **0.174** | **-3.976** | **<0.001** | **-1.032** | **-0.351** |
| **FO+AL** | **AL** | **-0.323** | **0.164** | **-1.974** | **0.048** | **-0.644** | **-0.002** |
| **AL** | **TF** | **-1.911** | **0.236** | **-8.101** | **<0.001** | **-2.373** | **-1.449** |
| **TF** | **TF** | **-2.244** | **0.249** | **-8.994** | **<0.001** | **-2.733** | **-1.755** |
| **FO** | **TF** | **-2.593** | **0.244** | **-10.629** | **<0.001** | **-3.071** | **-2.115** |
| **FO+AL** | **TF** | **-2.380** | **0.239** | **-9.967** | **<0.001** | **-2.847** | **-1.912** |

**Table S2F.** Model output from the best identified full model for Age-Specific Reproduction in $F_1$. Lineages are given as $P_0$-$F_1$.

| Model | Covariate | Estimate | SE | *z* value | *p* value |
|---|---|---|---|---|---|
| | **(Intercept)** | **3.750** | **0.165** | **22.744** | **<0.001** |
| | **TF-AL** | **-0.803** | **0.135** | **-5.939** | **<0.001** |
| | FO-AL | -0.036 | 0.124 | -0.286 | 0.775 |
| | FO+AL-AL | -0.032 | 0.120 | -0.266 | 0.790 |

| | | | | | |
|---|---|---|---|---|---|
| Conditional | **AL-TF** | **-3.015** | **0.212** | **-14.206** | **<0.001** |
| | **TF-TF** | **-7.689** | **0.366** | **-20.985** | **<0.001** |
| | **FO-TF** | **-3.572** | **0.217** | **-16.490** | **<0.001** |
| | **FO+AL-TF** | **-3.894** | **0.234** | **-16.665** | **<0.001** |
| | **Day2** | **-0.257** | **0.015** | **-17.169** | **<0.001** |
| | **Day** | **0.869** | **0.079** | **10.936** | **<0.001** |
| | TF-AL:Day2 | 0.034 | 0.019 | 1.758 | 0.079 |
| | FO-AL:Day2 | 0.028 | 0.019 | 1.437 | 0.151 |
| | **FO+AL-AL:Day2** | **0.056** | **0.018** | **3.003** | **0.003** |
| | **AL-TF:Day2** | **0.082** | **0.020** | **4.108** | **<0.001** |
| | **TF-TF:Day2** | **-0.109** | **0.023** | **-4.797** | **<0.001** |
| | **FO-TF:Day2** | **0.058** | **0.019** | **2.972** | **0.003** |
| | FO+AL-TF:Day2 | 0.047 | 0.020 | 2.389 | 0.017 |
| | TF-AL:Day | 0.139 | 0.110 | 1.261 | 0.207 |
| | FO-AL:Day | -0.074 | 0.107 | -0.696 | 0.487 |
| | FO+AL-AL:Day | -0.175 | 0.102 | -1.706 | 0.088 |
| | **AL-TF:Day** | **0.432** | **0.135** | **3.212** | **<0.001** |
| | **TF-TF:Day** | **2.528** | **0.178** | **14.210** | **<0.001** |
| | **FO-TF:Day** | **0.670** | **0.133** | **5.045** | **<0.001** |
| | **FO+AL-TF:Day** | **0.848** | **0.138** | **6.164** | **<0.001** |
| Zero-Inflation | **(Intercept)** | **-8.800** | **1.123** | **-7.834** | **<0.001** |
| | TF-AL | 0.540 | 0.307 | 1.757 | 0.079 |

| | | | | | |
|---|---|---|---|---|---|
| | FO-AL | -0.308 | 0.381 | -0.807 | 0.420 |
| | FO+AL-AL | -0.617 | 0.427 | -1.445 | 0.149 |
| | AL-TF | -1.122 | 0.665 | -1.689 | 0.091 |
| | **TF-TF** | **-1.672** | **0.745** | **-2.243** | **0.025** |
| | FO-TF | -4.121 | 4.709 | -0.875 | 0.382 |
| | FO+AL-TF | -1.227 | 0.676 | -1.814 | 0.070 |
| | **Day** | **2.631** | **0.495** | **5.313** | **<0.001** |
| | **Day2** | **-0.243** | **0.051** | **-4.752** | **<0.001** |

**Table S2G.** Model output from the best identified partial model for Age-Specific Reproduction in $F_1$ in temporary fasting conditions. Lineages are given as $P_0$-$F_1$.

| Model | Covariate | Estimate | SE | *z* value | *p* value |
|---|---|---|---|---|---|
| | (Intercept) | 0.267 | 0.328 | 0.815 | 0.415 |
| | **TF-TF** | **-4.981** | **0.405** | **-12.307** | **<0.001** |
| | **FO-TF** | **-0.581** | **0.278** | **-2.094** | **0.036** |
| | **FO+AL-TF** | **-0.939** | **0.290** | **-3.233** | **0.001** |
| | **Day2** | **-0.183** | **0.014** | **-13.267** | **<0.001** |
| | **Day** | **1.425** | **0.110** | **12.936** | **<0.001** |
| | **TF-TF:Day2** | **-0.199** | **0.022** | **-9.055** | **<0.001** |
| Conditional | FO-TF:Day2 | -0.023 | 0.019 | -1.238 | 0.216 |
| | FO+Al-TF:Day2 | -0.031 | 0.019 | -1.644 | 0.100 |
| | **TF-TF:Day** | **2.178** | **0.192** | **11.354** | **<0.001** |
| | FO-TF:Day | 0.243 | 0.151 | 1.610 | 0.107 |

| | | | | | |
|---|---|---|---|---|---|
| | **FO+Al-TF:Day** | **0.426** | **0.154** | **2.768** | **0.006** |
| Zero-Inflation | **(Intercept)** | **-50.161** | **14.569** | **-3.443** | **0.001** |
| | TF-TF | -0.993 | 0.626 | -1.585 | 0.113 |
| | FO-TF | -0.196 | 0.607 | -0.322 | 0.747 |
| | FO+AL-TF | 0.301 | 0.519 | 0.579 | 0.563 |
| | **Day** | **14.196** | **4.442** | **3.196** | **0.001** |
| | **Day2** | **-1.016** | **0.337** | **-3.012** | **0.003** |

**Table S2H.** Model output from the best identified partial model for Age-Specific Reproduction in $F_1$ in *ad libitum* conditions. Lineages are given as $P_0$-$F_1$.

| Model | Covariate | Estimate | SE | *z* value | *p* value |
|---|---|---|---|---|---|
| Conditional | **(Intercept)** | **3.905** | **0.137** | **28.552** | **<0.001** |
| | **TF-AL** | **-0.811** | **0.149** | **-5.451** | **<0.001** |
| | FO-AL | -0.062 | 0.137 | -0.448 | 0.654 |
| | FO+AL-AL | -0.088 | 0.133 | -0.662 | 0.508 |
| | **Day2** | **-0.232** | **0.015** | **-15.251** | **<0.001** |
| | **Day** | **0.732** | **0.084** | **8.678** | **<0.001** |
| | TF-AL:Day2 | 0.026 | 0.020 | 1.337 | 0.181 |
| | FO-AL:Day2 | 0.026 | 0.020 | 1.322 | 0.186 |
| | **FO+AL-AL:Day2** | **0.052** | **0.018** | **2.846** | **0.004** |
| | TF-AL:Day | 0.160 | 0.117 | 1.373 | 0.170 |
| | FO-AL:Day | -0.051 | 0.113 | -0.455 | 0.649 |
| | FO+AL-AL:Day | -0.131 | 0.107 | -1.218 | 0.223 |

**Table S2I.** Model output from the best identified full model for Total Reproduction in $F_1$. Lineages are given as $P_0$-$F_1$.

| Model | Covariate | Estimate | SE | *z* value | *p* value |
|---|---|---|---|---|---|
| | **(Intercept)** | **5.504** | **0.141** | **38.990** | **<0.001** |
| | **TF-AL** | **-0.162** | **0.050** | **-3.210** | **0.001** |
| | FO-AL | -0.022 | 0.046 | -0.470 | 0.638 |
| | FO+AL-AL | -0.050 | 0.045 | -1.120 | 0.261 |
| | **AL-TF** | **-1.391** | **0.085** | **-16.310** | **<0.001** |
| Conditional | **TF-TF** | **-0.745** | **0.076** | **-9.770** | **<0.001** |
| | **FO-TF** | **-1.481** | **0.079** | **-18.750** | **<0.001** |
| | **FO+AL-TF** | **-1.134** | **0.082** | **-13.780** | **<0.001** |
| Zero-Inflation | **(Intercept)** | **-5.2872** | **0.5851** | **-9.036** | **<0.001** |

**Table S2J.** Model output from the best identified partial model for Total Reproduction in $F_1$ in temporary fasting conditions. Lineages are given as $P_0$-$F_1$.

| Model | Covariate | Estimate | SE | *z* value | *p* value |
|---|---|---|---|---|---|
| | **(Intercept)** | **4.124** | **0.431** | **9.575** | **<0.001** |
| | **TF-TF** | **0.448** | **0.148** | **3.019** | **0.003** |
| Conditional | FO-TF | -0.095 | 0.124 | -0.768 | 0.442 |
| | FO+AL-TF | 0.232 | 0.134 | 1.731 | 0.083 |

**Table S2K.** Model output from the best identified partial model for Total Reproduction in $F_1$ in *ad libitum* conditions. Lineages are given as $P_0$-$F_1$.

| Model | Covariate | Estimate | SE | *z* value | *p* value |
|---|---|---|---|---|---|
| | **(Intercept)** | **5.495** | **0.102** | **54.110** | **<0.001** |

| | | | | | |
|---|---|---|---|---|---|
| Conditional | **TF-AL** | **-0.129** | **0.036** | **-3.590** | **<0.001** |
| | FO-AL | -0.012 | 0.032 | -0.370 | 0.713 |
| | FO+AL-AL | -0.045 | 0.031 | -1.460 | 0.144 |
| **Zero-Inflation** | **(Intercept)** | **-5.900** | **1.001** | **-5.892** | **<0.001** |

**Table S2L.** Model output from the full mixed effect model for individual fitness (λ) in $F_1$.

| Covariate | Estimate | SE | *z* value | *p* value |
|---|---|---|---|---|
| **(Intercept)** | **4.436** | **0.116** | **38.410** | **<0.001** |
| **TF-AL** | **-0.640** | **0.066** | **-9.670** | **<0.001** |
| FO-AL | -0.076 | 0.060 | -1.270 | 0.203 |
| **FO+AL-TF** | **-0.238** | **0.058** | **-4.090** | **0.000** |
| **AL-TF** | **-2.252** | **0.062** | **-36.420** | **<0.001** |
| **TF-TF** | **-2.433** | **0.080** | **-30.430** | **<0.001** |
| **FO-TF** | **-2.278** | **0.063** | **-36.120** | **<0.001** |
| **FO+AL-TF** | **-2.243** | **0.070** | **-32.080** | **<0.001** |

**Table S2M.** Model output from the full mixed effect model for individual fitness (λ) in $F_1$ in temporary fasting conditions.

| Covariate | Estimate | SE | *z* value | *p* value |
|---|---|---|---|---|
| **(Intercept)** | **2.193** | **0.089** | **24.541** | **<0.001** |
| TF-TF | -0.081 | 0.078 | -1.037 | 0.300 |
| FO-TF | -0.019 | 0.062 | -0.306 | 0.759 |
| FO+AL-TF | 0.003 | 0.067 | 0.050 | 0.960 |

**Table S2N.** Model output from the full mixed effect model for individual fitness ($\lambda$) in $F_1$ in *ad libitum* conditions.

| Covariate | Estimate | SE | *z* value | *p* value |
|---|---|---|---|---|
| (Intercept) | 4.432 | 0.147 | 30.114 | **<0.001** |
| TF-AL | -0.660 | 0.0720 | -9.131 | **<0.001** |
| FO-AL | -0.0770 | 0.0603 | -1.215 | 0.224 |
| FO+AL-AL | -0.243 | 0.0620 | -3.923 | **<0.001** |

**Table S2O.** Full model output from the Mixed Effects Cox Proportional Hazard Model for survival in $F_2$.

| $P_0$ | $F_1$ | $F_2$ | Estimate | SE | *z* value | *p* value | Low 95% CI | High 95% CI |
|---|---|---|---|---|---|---|---|---|
| **TF** | **AL** | **AL** | **1.211** | **0.212** | **5.705** | **<0.001** | **0.795** | **1.627** |
| **FO** | **AL** | **AL** | **0.972** | **0.205** | **4.731** | **<0.001** | **0.569** | **1.374** |
| FO+AL | AL | AL | 0.185 | 0.212 | 0.872 | 0.383 | -0.230 | 0.599 |
| **AL** | **AL** | **TF** | **-2.566** | **0.297** | **-8.646** | **<0.001** | **-3.148** | **-1.985** |
| **TF** | **AL** | **TF** | **-1.548** | **0.268** | **-5.782** | **<0.001** | **-2.073** | **-1.023** |
| **TF** | **TF** | **TF** | **-0.859** | **0.280** | **-3.068** | **0.002** | **-1.408** | **-0.310** |
| **FO** | **TF** | **TF** | **-1.001** | **0.306** | **-3.276** | **0.001** | **-1.600** | **-0.402** |

**Table S2P.** Model output from the best identified full model for Age-Specific Reproduction in $F_2$. Lineages are given as $P_0$-$F_1$-$F_2$.

| Model | Covariate | Estimate | SE | *z* value | *p* value |
|---|---|---|---|---|---|
| | **(Intercept)** | 3.642 | 0.105 | 34.770 | **<0.001** |
| | **TF-AL-AL** | 0.336 | 0.130 | 2.590 | **0.0095** |

| | | | | | |
|---|---|---|---|---|---|
| | FO-AL-AL | 0.188 | 0.134 | 1.410 | 0.159 |
| | FO+AL-AL-AL | 0.387 | 0.133 | 2.920 | 0.004 |
| Conditional | **TF-AL-AL** | **-3.361** | **0.258** | **-13.040** | **<0.001** |
| | **TF-AL-TF** | **-3.371** | **0.249** | **-13.540** | **<0.001** |
| | **TF-TF-TF** | **-4.293** | **0.308** | **-13.920** | **<0.001** |
| | **FO-TF-TF** | **-4.392** | **0.365** | **-12.020** | **<0.001** |
| | **Day2** | **-0.274** | **0.016** | **-16.710** | **<0.001** |
| | **Day** | **1.016** | **0.088** | **11.490** | **<0.001** |
| | **TF-AL-AL:Day2** | **0.102** | **0.020** | **5.220** | **<0.001** |
| | FO-AL-AL:Day2 | 0.054 | 0.021 | 2.570 | 0.0103 |
| | **FO+AL-AL-AL:Day2** | **0.084** | **0.021** | **4.070** | **<0.001** |
| | **TF-AL-AL:Day2** | **0.095** | **0.021** | **4.510** | **<0.001** |
| | TF-AL-TF:Day2 | 0.070 | 0.021 | 3.290 | 0.001 |
| | TF-TF-TF:Day2 | 0.018 | 0.023 | 0.790 | 0.429 |
| | FO-TF-TF:Day2 | 0.024 | 0.026 | 0.950 | 0.343 |
| | **TF-AL-AL:Day** | **-0.432** | **0.110** | **-3.940** | **<0.001** |
| | FO-AL-AL:Day | -0.222 | 0.115 | -1.930 | 0.0540 |
| | **FO+AL-AL-AL:Day** | **-0.412** | **0.114** | **-3.610** | **<0.001** |
| | AL-AL-TF:Day | 0.487 | 0.149 | 3.270 | 0.00108 |
| | **TF-AL-TF:Day** | **0.642** | **0.148** | **4.340** | **<0.001** |
| | **TF-TF-TF:Day** | **1.105** | **0.169** | **6.550** | **<0.001** |
| | **FO-TF-TF:Day** | **1.078** | **0.198** | **5.460** | **<0.001** |

| | | | | | |
|---|---|---|---|---|---|
| Zero-Inflation | **(Intercept)** | **-15.903** | **1.797** | **-8.848** | **<0.001** |
| | **TF-AL-AL** | **1.373** | **0.400** | **3.433** | **<0.001** |
| | FO-AL-AL | 0.934 | 0.419 | 2.227 | 0.0259 |
| | FO+AL-AL-AL | 0.831 | 0.425 | 1.958 | 0.0502 |
| | AL-AL-TF | -0.743 | 0.716 | -1.038 | 0.299 |
| | TF-AL-TF | -0.885 | 0.725 | -1.222 | 0.222 |
| | TF-TF-TF | -1.607 | 1.038 | -1.549 | 0.121 |
| | FO-TF-TF | -17.702 | 2818.174 | -0.006 | 0.995 |
| | **Day** | **5.226** | **0.747** | **6.992** | **<0.001** |
| | **Day2** | **-0.488** | **0.077** | **-6.346** | **<0.001** |

**Table S2Q.** Model output from the best identified partial model for Age-Specific Reproduction in $F_2$ in temporary fasting conditions. Lineages are given as $P_0$-$F_1$-$F_2$.

| Model | Covariate | Estimate | SE | *z* value | *p* value |
|---|---|---|---|---|---|
| Conditional | (Intercept) | 0.146 | 0.250 | 0.584 | 0.559 |
| | TF-AL-TF | 0.030 | 0.326 | 0.092 | 0.927 |
| | **TF-TF-TF** | **-1.005** | **0.384** | **-2.614** | **0.009** |
| | FO-TF-TF | -0.742 | 0.432 | -1.716 | 0.086 |
| | **Day2** | **-0.159** | **0.014** | **-11.267** | **<0.001** |
| | **Day** | **1.444** | **0.124** | **11.638** | **<0.001** |
| | TF-AL-TF:Day2 | -0.027 | 0.020 | -1.350 | 0.177 |
| | **TF-TF-TF:Day2** | **-0.089** | **0.023** | **-3.880** | **<0.001** |
| | FO-TF-TF:Day2 | -0.054 | 0.026 | -2.029 | 0.042 |

| | | | | | |
|---|---|---|---|---|---|
| | TF-AL-TF:Day | 0.144 | 0.170 | 0.847 | 0.397 |
| | **TF-TF-TF:Day** | **0.679** | **0.195** | **3.475** | **0.001** |
| | FO-TF-TF:Day | 0.434 | 0.225 | 1.928 | 0.054 |
| Zero-Inflation | **(Intercept)** | **-10.883** | **1.241** | **-8.768** | **<0.001** |
| | TF-AL-TF | -0.191 | 0.444 | -0.431 | 0.667 |
| | TF-TF-TF | -0.532 | 0.520 | -1.024 | 0.306 |
| | FO-TF-TF | 0.061 | 0.480 | 0.127 | 0.899 |
| | **Day** | **1.455** | **0.175** | **8.323** | **<0.001** |

**Table S2R.** Model output from the best identified partial model for Age-Specific Reproduction in $F_2$ in *ad libitum* conditions. Lineages are given as $P_0$-$F_1$-$F_2$.

| Model | Covariate | Estimate | SE | *z* value | *p* value |
|---|---|---|---|---|---|
| Conditional | **(Intercept)** | **3.579** | **0.109** | **32.840** | **<0.001** |
| | **TF-AL-AL** | **0.349** | **0.131** | **2.660** | **0.008** |
| | FO-AL-AL | 0.177 | 0.135 | 1.310 | 0.191 |
| | **FO+AL-AL-AL** | **0.339** | **0.135** | **2.510** | **0.012** |
| | **Day2** | **-0.277** | **0.016** | **-16.870** | **<0.001** |
| | **Day** | **1.051** | **0.088** | **11.930** | **<0.001** |
| | **TF-AL-AL:Day2** | **0.103** | **0.019** | **5.280** | **<0.001** |
| | **FO-AL-AL:Day2** | **0.051** | **0.021** | **2.460** | **0.014** |
| | **FO+AL-AL-AL:Day2** | **0.072** | **0.021** | **3.450** | **0.001** |
| | **TF-AL-AL:Day** | **-0.434** | **0.108** | **-4.000** | **<0.001** |
| | FO-AL-AL:Day | -0.206 | 0.113 | -1.820 | 0.069 |

| | | | | | |
|---|---|---|---|---|---|
| | **FO+AL-AL-AL:Day** | **-0.355** | **0.114** | **-3.120** | **0.002** |
| Zero-Inflation | **(Intercept)** | **-16.138** | **2.637** | **-6.119** | **<0.001** |
| | TF-AL-AL | -1.463 | 2.164 | -0.676 | 0.499 |
| | FO-AL-AL | -0.350 | 2.234 | -0.157 | 0.875 |
| | FO+AL-AL-AL | 2.355 | 2.354 | 1.000 | 0.317 |
| | **Day** | **5.687** | **0.953** | **5.969** | **<0.001** |
| | **Day2** | **-0.585** | **0.093** | **-6.318** | **<0.001** |
| | TF-AL-AL:Day | 0.649 | 0.496 | 1.310 | 0.190 |
| | FO-AL-AL:Day | 0.309 | 0.516 | 0.600 | 0.548 |
| | FO+AL-AL-AL:Day | -0.355 | 0.570 | -0.623 | 0.533 |

**Table S2S.** Model output from the best identified full model for Total Reproduction in $F_2$. Lineages are given as $P_0$-$F_1$-$F_2$.

| Model | Covariate | Estimate | SE | *z* value | *p* value |
|---|---|---|---|---|---|
| Conditional | **(Intercept)** | **5.625** | **0.055** | **102.240** | **<0.001** |
| | TF-AL-AL | 0.023 | 0.045 | 0.510 | 0.607 |
| | FO-AL-AL | 0.043 | 0.045 | 0.960 | 0.339 |
| | FOAL-AL-AL | 0.012 | 0.046 | 0.250 | 0.804 |
| | **AL-AL-TF** | **-1.223** | **0.082** | **-14.940** | **<0.001** |
| | **TF-AL-TF** | **-0.973** | **0.082** | **-11.820** | **<0.001** |
| | **TF-TF-TF** | **-0.718** | **0.079** | **-9.130** | **<0.001** |
| | **FO-TF-TF** | **-1.067** | **0.093** | **-11.460** | **<0.001** |
| Zero-Inflation | **(Intercept)** | **-5.361** | **0.709** | **-7.564** | **<0.001** |

**Table S2T.** Model output from the best identified partial model for Total Reproduction in $F_2$ in temporary fasting conditions. Lineages are given as $P_0$-$F_1$-$F_2$.

| Model | Covariate | Estimate | SE | *z* value | *p* value |
|---|---|---|---|---|---|
| | **(Intercept)** | **4.520** | **0.110** | **41.250** | **<0.001** |
| | TF-AL-TF | 0.190 | 0.149 | 1.270 | 0.204 |
| Conditional | **TF-TF-TF** | **0.392** | **0.165** | **2.370** | **0.018** |
| | FO-TF-TF | 0.126 | 0.174 | 0.720 | 0.470 |

**Table S2U.** Model output from the best identified partial model for Total Reproduction in $F_2$ in *ad libitum* conditions. Lineages are given as $P_0$-$F_1$-$F_2$.

| Model | Covariate | Estimate | SE | *z* value | *p* value |
|---|---|---|---|---|---|
| | **(Intercept)** | **5.594** | **0.063** | **89.110** | **<0.001** |
| | **TF-AL-AL** | 0.032 | 0.024 | 1.320 | 0.187 |
| Conditional | FO-AL-AL | 0.039 | 0.024 | 1.640 | 0.102 |
| | **FO+AL-AL-AL** | 0.015 | 0.025 | 0.580 | 0.564 |
| Zero-Inflation | **(Intercept)** | **-4.977** | **0.710** | **-7.014** | **<0.001** |

**Table S2V.** Model output from the full mixed effect model for individual fitness (λ) in $F_2$.

| Covariate | Estimate | SE | *z* value | *p* value |
|---|---|---|---|---|
| **(Intercept)** | **4.457** | **0.079** | **56.430** | **<0.001** |
| TF-AL-AL | -0.039 | 0.059 | -0.660 | 0.512 |
| FO-AL-AL | -0.037 | 0.060 | -0.620 | 0.537 |
| FO+AL-AL-AL | -0.062 | 0.064 | -0.970 | 0.332 |
| **AL-AL-TF** | **-2.163** | **0.051** | **-42.130** | **<0.001** |

| | | | | |
|---|---|---|---|---|
| **TF-AL-TF** | **-2.096** | **0.068** | **-30.990** | **<0.001** |
| **TF-TF-TF** | **-1.956** | **0.103** | **-18.950** | **<0.001** |
| **FO-TF-TF** | **-2.061** | **0.094** | **-21.940** | **<0.001** |

**Table S2W.** Model output from the full mixed effect model for individual fitness (λ) in $F_2$ in temporary fasting conditions.

| Covariate | Estimate | SE | *z* value | *p* value |
|---|---|---|---|---|
| **(Intercept)** | **2.292** | **0.036** | **64.530** | **<0.001** |
| TF-AL-TF | 0.066 | 0.052 | 1.270 | 0.203 |
| **TF-TF-TF** | **0.185** | **0.061** | **3.020** | **0.003** |
| **FO-TF-TF** | 0.070 | 0.061 | 1.140 | 0.254 |

**Table S2X.** Model output from the full mixed effect model for individual fitness (λ) in $F_2$ in *ad libitum* conditions.

| Covariate | Estimate | SE | *z* value | *p* value |
|---|---|---|---|---|
| **(Intercept)** | **4.460** | **0.104** | **42.710** | **<0.001** |
| TF-AL-AL | -0.042 | 0.065 | -0.650 | 0.515 |
| FO-AL-AL | -0.032 | 0.065 | -0.500 | 0.619 |
| FO+AL-AL-AL | -0.070 | 0.072 | -0.970 | 0.332 |

**Table S2Y.** Full model output from the Mixed Effects Cox Proportional Hazard Model for survival in $F_3$.

| $P_0$ | $F_1$ | $F_2$ | $F_3$ | Estimate | SE | *z* value | *p* value | Low 95% CI | High 95% CI |
|---|---|---|---|---|---|---|---|---|---|
| **TF** | **AL** | **AL** | **AL** | **1.308** | **0.192** | **6.827** | **<0.001** | **0.933** | **1.684** |
| **FO** | **AL** | **AL** | **AL** | **1.314** | **0.192** | **6.828** | **<0.001** | **0.937** | **1.692** |

| | | | | | | | | | |
|---|---|---|---|---|---|---|---|---|---|
| FO+AL | AL | AL | AL | -0.137 | 0.182 | -0.756 | 0.450 | -0.494 | 0.219 |
| **AL** | **AL** | **AL** | **TF** | **-1.928** | **0.259** | **-7.445** | **<0.001** | **-2.435** | **-1.420** |
| **TF** | **AL** | **AL** | **TF** | **-1.010** | **0.232** | **-4.359** | **<0.001** | **-1.465** | **-0.556** |
| TF | TF | TF | TF | 0.395 | 0.256 | 1.541 | 0.123 | -0.107 | 0.898 |
| FO | TF | TF | TF | -0.085 | 0.287 | -0.297 | 0.766 | -0.647 | 0.477 |

**Table S2Z.** Model output from the best identified full model for Age-Specific Reproduction in $F_3$. Lineages are given as $P_0$-$F_1$-$F_2$-$F_3$.

| Model | Covariate | Estimate | SE | *z* value | *p* value |
|---|---|---|---|---|---|
| Conditional | **(Intercept)** | **3.367** | **0.107** | **31.391** | **<0.001** |
| | TF-AL-AL-AL | 0.202 | 0.129 | 1.560 | 0.119 |
| | FO-AL-AL-AL | 0.242 | 0.127 | 1.904 | 0.057 |
| | FO+AL-AL-AL-AL | 0.095 | 0.133 | 0.714 | 0.475 |
| | **AL-AL-AL-TF** | **-3.781** | **0.309** | **-12.243** | **<0.001** |
| | **TF-AL-AL-TF** | **-4.691** | **0.309** | **-15.181** | **<0.001** |
| | **TF-TF-TF-TF** | **-5.830** | **0.389** | **-14.971** | **<0.001** |
| | **FO-TF-TF-TF** | **-5.757** | **0.475** | **-12.120** | **<0.001** |
| | **Day2** | **-0.291** | **0.014** | **-20.676** | **<0.001** |
| | **Day** | **1.185** | **0.078** | **15.103** | **<0.001** |
| | **TF-AL-AL-AL:Day2** | **0.045** | **0.018** | **2.493** | **0.013** |
| | **FO-AL-AL-AL:Day2** | **0.060** | **0.018** | **3.402** | **0.001** |
| | FO+AL-AL-AL-AL:Day2 | 0.006 | 0.019 | 0.325 | 0.745 |
| | **AL-AL-AL-TF:Day2** | **0.071** | **0.021** | **3.373** | **0.001** |

| | | | | | |
|---|---|---|---|---|---|
| | TF-AL-AL-TF:Day2 | 0.033 | 0.020 | 1.630 | 0.103 |
| | TF-TF-TF-TF:Day2 | 0.031 | 0.021 | 1.438 | 0.150 |
| | FO-TF-TF-TF:Day2 | 0.000 | 0.026 | -0.015 | 0.988 |
| | **TF-AL-AL-AL:Day** | **-0.247** | **0.104** | **-2.383** | **0.017** |
| | **FO-AL-AL-AL:Day** | **-0.290** | **0.101** | **-2.874** | **0.004** |
| | FO+AL-AL-AL-AL:Day | -0.073 | 0.108 | -0.679 | 0.497 |
| | **AL-AL-AL-TF:Day** | **0.700** | **0.164** | **4.273** | **<0.001** |
| | **TF-AL-AL-TF:Day** | **1.118** | **0.156** | **7.176** | **<0.001** |
| | **TF-TF-TF-TF:Day** | **1.417** | **0.179** | **7.934** | **<0.001** |
| | **FO-TF-TF-TF:Day** | **1.525** | **0.221** | **6.902** | **<0.001** |

**Table S2AA.** Model output from the best identified partial model for Age-Specific Reproduction in $F_3$ in temporary fasting conditions. Lineages are given as $P_0$-$F_1$-$F_2$-$F_3$.

| Model | Covariate | Estimate | SE | *z* value | *p* value |
|---|---|---|---|---|---|
| | **(Intercept)** | **-0.811** | **0.312** | **-2.599** | **0.009** |
| | TF-AL-AL-TF | -0.607 | 0.407 | -1.492 | 0.136 |
| | **TF-TF-TF-TF** | **-2.318** | **0.466** | **-4.970** | **<0.001** |
| | **FO-TF-TF-TF** | **-2.154** | **0.560** | **-3.849** | **<0.001** |
| | **Day2** | **-0.222** | **0.018** | **-12.313** | **<0.001** |
| | **Day** | **1.979** | **0.155** | **12.728** | **<0.001** |
| Conditional | TF-AL-AL-TF:Day2 | -0.010 | 0.023 | -0.434 | 0.664 |
| | TF-TF-TF-TF:Day2 | -0.064 | 0.023 | -2.722 | 0.006 |
| | FO-TF-TF-TF:Day2 | -0.087 | 0.030 | -2.959 | 0.003 |

| | | | | | |
|---|---|---|---|---|---|
| | TF-AL-AL-TF:Day | 0.215 | 0.199 | 1.079 | 0.280 |
| | **TF-TF-TF-TF:Day** | **0.874** | **0.214** | **4.084** | **<0.001** |
| | **FO-TF-TF-TF:Day** | **0.934** | **0.263** | **3.548** | **<0.001** |
| Zero-Inflation | **(Intercept)** | **-11.184** | **1.873** | **-5.972** | **<0.001** |
| | TF-AL-AL-TF | 0.451 | 0.508 | 0.887 | 0.375 |
| | TF-TF-TF-TF | -23.561 | 29875.966 | -0.001 | 0.999 |
| | FO-TF-TF-TF | -1.463 | 1.257 | -1.164 | 0.244 |
| | **Day** | **1.402** | **0.256** | **5.475** | **<0.001** |

**Table S2AB.** Model output from the best identified partial model for Age-Specific Reproduction in $F_3$ in *ad libitum* conditions. Lineages are given as $P_0$-$F_1$-$F_2$-$F_3$.

| Model | Covariate | Estimate | SE | *z* value | *p* value |
|---|---|---|---|---|---|
| Conditional | **(Intercept)** | **3.413** | **0.107** | **31.850** | **<0.001** |
| | TF-AL-AL-AL | 0.153 | 0.124 | 1.240 | 0.215 |
| | FO-AL-AL-AL | 0.200 | 0.121 | 1.650 | 0.099 |
| | FO+AL-AL-AL-AL | -0.002 | 0.127 | -0.020 | 0.986 |
| | **Day2** | **-0.292** | **0.014** | **-20.850** | **<0.001** |
| | **Day** | **1.178** | **0.077** | **15.280** | **<0.001** |
| | **TF-AL-AL-AL:Day2** | **0.042** | **0.018** | **2.380** | **0.017** |
| | **FO-AL-AL-AL:Day2** | **0.054** | **0.017** | **3.170** | **0.002** |
| | FO+AL-AL-AL-AL:Day2 | -0.006 | 0.019 | -0.300 | 0.767 |
| | **TF-AL-AL-AL:Day** | **-0.221** | **0.101** | **-2.200** | **0.028** |
| | **FO-AL-AL-AL:Day** | **-0.258** | **0.098** | **-2.640** | **0.008** |

| | | | | | |
|---|---|---|---|---|---|
| | FO+AL-AL-AL-AL:Day | 0.011 | 0.104 | 0.100 | 0.917 |
| Zero-Inflation | **(Intercept)** | **-4.131** | **0.228** | **-18.160** | **<0.001** |

**Table S2AC.** Model output from the best identified full model for Total Reproduction in $F_3$. Lineages are given as $P_0$-$F_1$-$F_2$-$F_3$.

| Model | Covariate | Estimate | SE | *z* value | *p* value |
|---|---|---|---|---|---|
| Conditional | (Intercept) | 5.666 | 0.052 | 109.070 | **<0.001** |
| | TF-AL-AL-AL | -0.067 | 0.044 | -1.510 | 0.130 |
| | FO-AL-AL-AL | -0.044 | 0.044 | -1.000 | 0.318 |
| | FO+AL-AL-AL-AL | -0.015 | 0.045 | -0.330 | 0.744 |
| | **AL-AL-AL-TF** | **-1.061** | **0.083** | **-12.780** | **<0.001** |
| | **TF-AL-AL-TF** | **-0.788** | **0.073** | **-10.760** | **<0.001** |
| | **TF-TF-TF-TF** | **-0.637** | **0.078** | **-8.150** | **<0.001** |
| | **FO-TF-TF-TF** | **-0.817** | **0.107** | **-7.630** | **<0.001** |
| Zero-Inflation | (Intercept) | -19.287 | 1817.670 | -0.011 | 0.992 |
| | TF-AL-AL-AL | -4.340 | 15700.221 | 0.000 | 1.000 |
| | FO-AL-AL-AL | -4.299 | 14998.436 | 0.000 | 1.000 |
| | FO+AL-AL-AL-AL | 15.053 | 1817.671 | 0.008 | 0.993 |
| | AL-AL-AL-TF | -4.489 | 25785.964 | 0.000 | 1.000 |
| | TF-AL-AL-TF | -4.512 | 25312.863 | 0.000 | 1.000 |
| | TF-TF-TF-TF | 17.128 | 1817.670 | 0.009 | 0.992 |
| | FO-TF-TF-TF | 17.272 | 1817.670 | 0.010 | 0.992 |

**Table S2AD.** Model output from the best identified partial model for Total Reproduction in $F_3$ in temporary fasting conditions. Lineages are given as $P_0$-$F_1$-$F_2$-$F_3$.

| Model | Covariate | Estimate | SE | *z* value | *p* value |
|---|---|---|---|---|---|
| Conditional | **(Intercept)** | **4.634** | **0.133** | **34.950** | **<0.001** |
| | TF-AL-AL-TF | 0.247 | 0.143 | 1.730 | 0.083 |
| | **TF-TF-TF-TF** | **0.352** | **0.152** | **2.320** | **0.021** |
| | FO-TF-TF-TF | 0.251 | 0.187 | 1.350 | 0.178 |
| Zero-Inflated | (Intercept) | -20.227 | 4361.236 | -0.005 | 0.996 |
| | TF-AL-AL-TF | -4.756 | 45826.925 | 0.000 | 1.000 |
| | **TF-TF-TF-TF** | 18.067 | 4361.236 | 0.004 | 0.997 |
| | FO-TF-TF-TF | 18.212 | 4361.236 | 0.004 | 0.997 |

**Table S2AE.** Model output from the best identified partial model for Total Reproduction in $F_3$ in *ad libitum* conditions. Lineages are given as $P_0$-$F_1$-$F_2$-$F_3$.

| Model | Covariate | Estimate | SE | *z* value | *p* value |
|---|---|---|---|---|---|
| Conditional | **(Intercept)** | **5.644** | **0.070** | **80.400** | **<0.001** |
| | **TF-AL-AL-AL** | **-0.060** | **0.028** | **-2.150** | **0.032** |
| | FO-AL-AL-AL | -0.034 | 0.028 | -1.220 | 0.224 |
| | FO+AL-AL-AL-AL | -0.014 | 0.029 | -0.500 | 0.617 |
| Zero-Inflation | **(Intercept)** | **-5.687** | **1.002** | **-5.677** | **<0.001** |

**Table S2AF.** Model output from the full mixed effect model for individual fitness (λ) in $F_3$.

| Covariate | Estimate | SE | *z* value | *p* value |
|---|---|---|---|---|
| **(Intercept)** | **4.419** | **0.122** | **36.260** | **<0.001** |
| TF-AL-AL-AL | -0.050 | 0.072 | -0.690 | 0.489 |

| | | | | |
|---|---|---|---|---|
| FO-AL-AL-AL | -0.044 | 0.072 | -0.610 | 0.542 |
| FO+AL-AL-AL-AL | -0.035 | 0.074 | -0.480 | 0.630 |
| **AL-AL-AL-TF** | **-2.010** | **0.092** | **-21.900** | **<0.001** |
| **TF-AL-AL-TF** | **-2.071** | **0.091** | **-22.690** | **<0.001** |
| **TF-TF-TF-TF** | **-2.326** | **0.097** | **-23.890** | **<0.001** |
| **FO-TF-TF-TF** | **-2.427** | **0.120** | **-20.300** | **<0.001** |

**Table S2AG.** Model output from the full mixed effect model for individual fitness ($\lambda$) in $F_3$ in temporary fasting conditions.

| Covariate | Estimate | SE | *z* value | *p* value |
|---|---|---|---|---|
| **(Intercept)** | **2.373** | **0.117** | **20.235** | **<0.001** |
| TF-AL-AL-TF | -0.033 | 0.123 | -0.267 | 0.789 |
| **TF-TF-TF-TF** | **-0.318** | **0.127** | **-2.509** | **0.012** |
| **FO-TF-TF-TF** | **-0.399** | **0.151** | **-2.637** | **0.008** |

**Table S2AH.** Model output from the full mixed effect model for individual fitness ($\lambda$) in $F_3$ in *ad libitum* conditions.

| Covariate | Estimate | SE | *z* value | *p* value |
|---|---|---|---|---|
| **(Intercept)** | **4.421** | **0.127** | **34.860** | **<0.001** |
| TF-AL-AL-AL | -0.053 | 0.069 | -0.760 | 0.444 |
| FO-AL-AL-AL | -0.049 | 0.068 | -0.720 | 0.472 |
| FO+AL-AL-AL-AL | -0.040 | 0.070 | -0.560 | 0.573 |
